# *Slc7a8* deletion is protective against diet-induced obesity and attenuates lipid accumulation in multiple organs

**DOI:** 10.1101/2021.10.25.465668

**Authors:** Reabetswe R. Pitere, Marlene B. van Heerden, Michael S. Pepper, Melvin A. Ambele

## Abstract

Adipogenesis, through adipocyte hyperplasia and/or hypertrophy, leads to increased adiposity, giving rise to obesity. A genome-wide transcriptome analysis of adipogenesis in human adipose derived stromal/stem cells identified SLC7A8 (Solute Carrier Family 7 Member 8) as a potential novel mediator. This study has investigated the role of SLC7A8 in adipose tissue biology using a mouse model of diet-induced obesity. *slc7a8* knockout (KO) and wildtype (WT) C57BL/6J mice were fed either a control diet (CD) or a high-fat diet (HFD) for 14 weeks. On HFD, both WT and KO (WTHFD and KOHFD) gained significantly more weight than their CD counterparts. However, KOHFD gained significantly less weight than WT HFD. KOHFD significantly reduced the level of glucose intolerance observed in WTHFD. KOHFD significantly reduced both adipocyte mass and hypertrophy in inguinal, mesenteric, perigonadal and brown adipose depots, with a corresponding decrease in macrophage infiltration. Additionally, KOHFD decreased lipid accumulation in the liver, heart, gastrocnemius muscle, lung, and kidney. This study demonstrates that targeting *SLC7A8* protects against diet-induced obesity by reducing lipid accumulation in multiple organs, and thereby has the potential to mitigate the development of obesity-associated comorbidities.

**Author summary:** The development of obesity can be attributed to adipocyte hypertrophy or hyperplasia leading to increased adiposity. The C57BL/6 mouse is an excellent model used to study metabolic syndromes often associated with obesity development. Mice fed on a high-fat diet are susceptible to weight gain leading to the development of obesity and its associated metabolic syndrome. Here, we report findings from targeting a novel human adipogenic gene (*SLC7A8*) in condition of obesity development using a mouse model of diet-induced obesity (DIO). The results indicate that deleting *slc7a8* in mice significantly protects against DIO and improves glucose metabolism. Also, deficiency in *slc7a8* was observed to significantly attenuates adipocyte hypertrophy in white and brown adipose tissues, and reduced lipid accumulation in many organs. Furthermore, inflammation was significantly reduced in adipose tissues and liver of *slc7a8* deficient mice in condition of DIO. Overall, results from this study shows that *slc7a8* is an important molecular regulator of obesity development and mediates its function by reducing lipid accumulation in multiple organs. Hence, SLC7A8 could serve as a potential therapeutic target to combat the development of obesity and other pathophysiological conditions associated with excess lipid accumulation in organs.

## Introduction

Obesity is characterised by an excess accumulation of adipose tissue when energy intake exceeds energy expenditure. The expansion of adipose tissue in obesity occurs either through adipocyte hyperplasia or hypertrophy. The result is dysfunctional adipose tissue mainly due to adipocyte hypertrophy, which leads to adverse metabolic consequences and chronic inflammation[1]. The distribution of adipose tissue in obesity plays an important role in the development of obesity-associated comorbidities. Accumulation of fat in the intra-abdominal depots (visceral depots) gives rise to insulin resistance and is also associated with an increased risk of cardiovascular diseases[2]. Subcutaneous white adipose tissue (WAT) is the most common adipose tissue in healthy lean individuals and serves as a metabolic sink for excess lipid storage[3]. Brown adipose tissue takes up fatty acid in circulation to generate heat, which helps to clear plasma triglycerides thereby reducing the accumulation of lipid at visceral depots[4]. In obesity, where the storage capacity of adipose tissue is exceeded either due to an inability to produce new adipocytes (limited hyperplasia) or to expand further (limited hypertrophy), excess fat is redistributed to peripheral organs such as the liver and skeletal muscle which increases the risk of metabolic syndromes such as hyperglycaemia, hyperinsulinemia, atherosclerosis, dyslipidemia and systemic inflammation[3, 5]. Hypertrophy in brown adipose tissue (BAT) may impair its function in acting as a sink for excess blood glucose and clearance of free fatty acids from circulation, thereby contributing to the development of insulin resistance and hyperlipidemia in obesity[3]. Therefore, mitigating adipocyte hypertrophy in both WAT and BAT depots is paramount to improving metabolic health.

Inflammation is a key consequence of adipose tissue expansion that occurs during weight gain and contributes to the development of chronic low-grade systemic inflammation seen in obesity. This expansion of adipose tissue is characterized by increased infiltration of immune cells, with a predominance (around 60%) of macrophages, in response to chemokines that are produced by hypertrophic adipocytes[6]. The majority are derived from circulating monocytes with a small proportion coming from the proliferation of adipose tissue resident macrophages[7]. Tissue resident macrophages present in normal or lean adipose tissue are of the M2 anti-inflammatory macrophage phenotype that express markers such as mannose receptor (CD206), and are thought to be responsible for maintaining tissue homeostasis[8]. Macrophage infiltration in adipose tissue appears as crown-like clusters which is believed to signify an immune response to dying or dead adipocytes[9]. These infiltrating macrophages undergo a phenotypic switch to a M1 pro-inflammatory phenotype[10].

In addressing obesity, several studies have suggested exploiting the process of fat cell formation (adipogenesis) to combat obesity development. These have led to several molecular determinants being described to play important role in adipogenesis[11]. Except for PPARɣ[12, 13], molecular determinants of adipogenesis have proven to be of limited clinical utility. Therefore, more research is needed to identify new molecular determinants of adipogenesis which could play a role in obesity development and serve potential therapeutic targets. We have previously undertaken an unbiased exploratory comprehensive transcriptomic analysis of human adipose-derived stromal/stem cells undergoing adipogenesis and identified several novel genes and transcription factors with possible role in this process [14, 15]. One of the novel genes identified was *SLC7A8* (Solute Carrier Family 7 Member 8), not previously described in the context of adipogenesis and/or obesity, that was significantly upregulated in the early phase of adipogenesis and declined significantly as the process progressed [14]. This could suggest a role for this gene in the early stages of adipogenesis as a potential driver of adiposity and consequently obesity. The aim of this study was to therefore investigate the functional role of the *SLC7A8* in weight gain/obesity development and lipid accumulation in various organs/tissues using a mouse model of diet induced obesity, as well as the macrophage infiltration profile in some of these tissues.

## Results

### Deficiency of *slc7a8* protects against diet-induced obesity

WT and KO mice significantly gained weight at 14 weeks on HFD compared to WTCD (p<0.05 to p<0.001) and KOCD (p<0.05 to p<0.001) (Figure 1A). No significant differences were observed between WT and KO on CD. Interestingly, KOHFD gained significantly (p<0.05 to p<0.001) less weight than WTHFD, which was evident from week 3 (Figure 1A). Significant weight gain in WTHFD was associated with significantly larger (p<0.001) iWAT, pWAT, mWAT, BAT and liver compared to WTCD and KOHFD. Only the pWAT of KOHFD was significantly larger than in KOCD14 (Figure 1D). WTHFD and KOHFD mice appear visibly larger in size when compared to their respective lean counterparts (Figure 1 supplementary). Food consumption was similar across the four groups under study, except that WTHFD had significantly greater food intake at 7 weeks when compared to KOHFD (p<0.01) and at 11 weeks when compared to WTCD (p<0.01) and KOHFD (p<0.001) (Figure 1B). Energy intake increased significantly between WTCD and WTHFD (p<0.01 at week 5 and p<0.001 from week 6 to week 14) and between KOCD and KOHFD (p<0.05 at week 3, p<0.01 at week 4, and p<0.001 from week 5 to 14) (Figure 1C). A significant difference (p<0.001) in cumulative caloric intake was observed between WTHFD and KOHFD from week 9 to week 14. No significant differences in calorie intake were seen between KOCD and WTCD.

**Figure 1:**
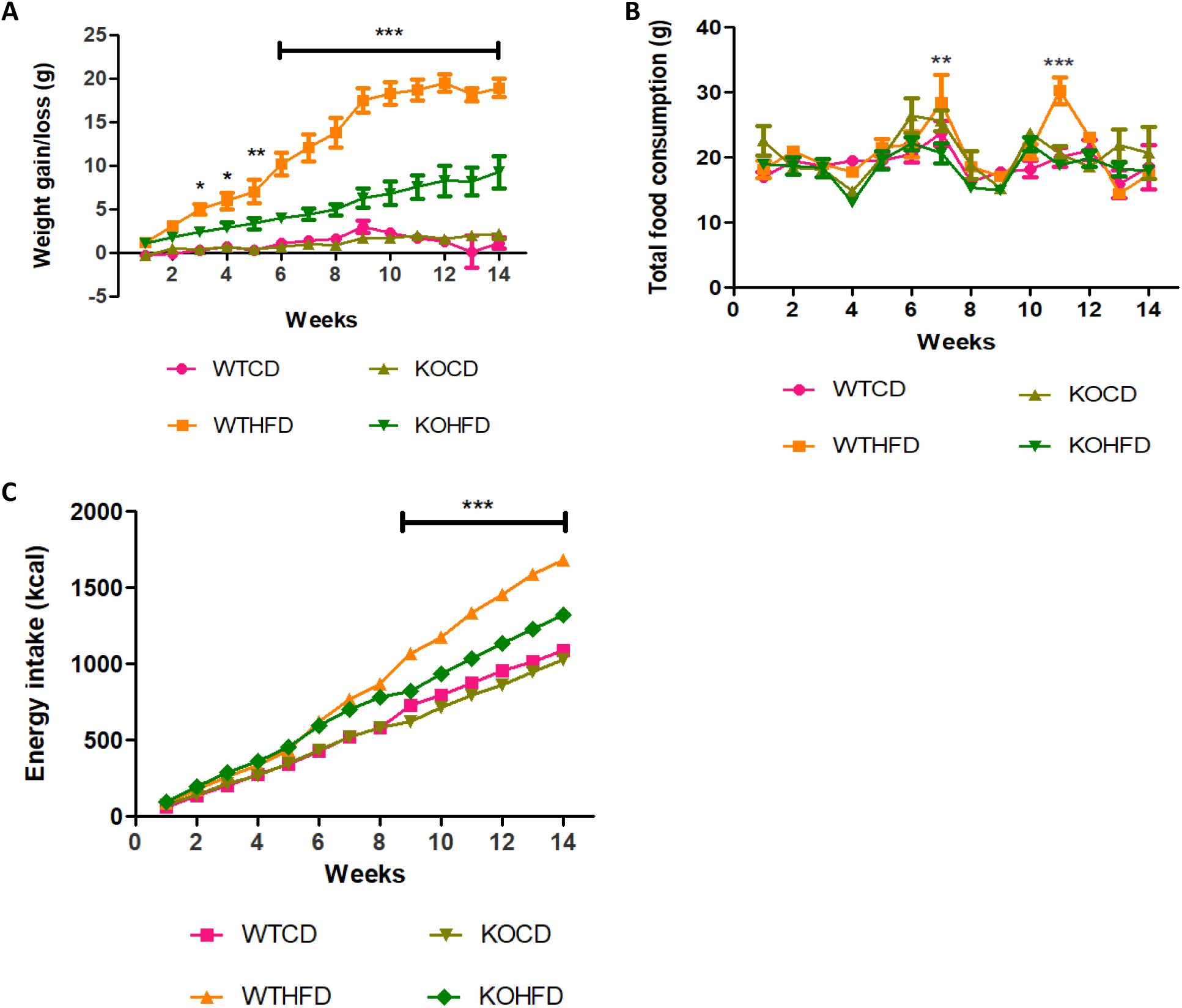
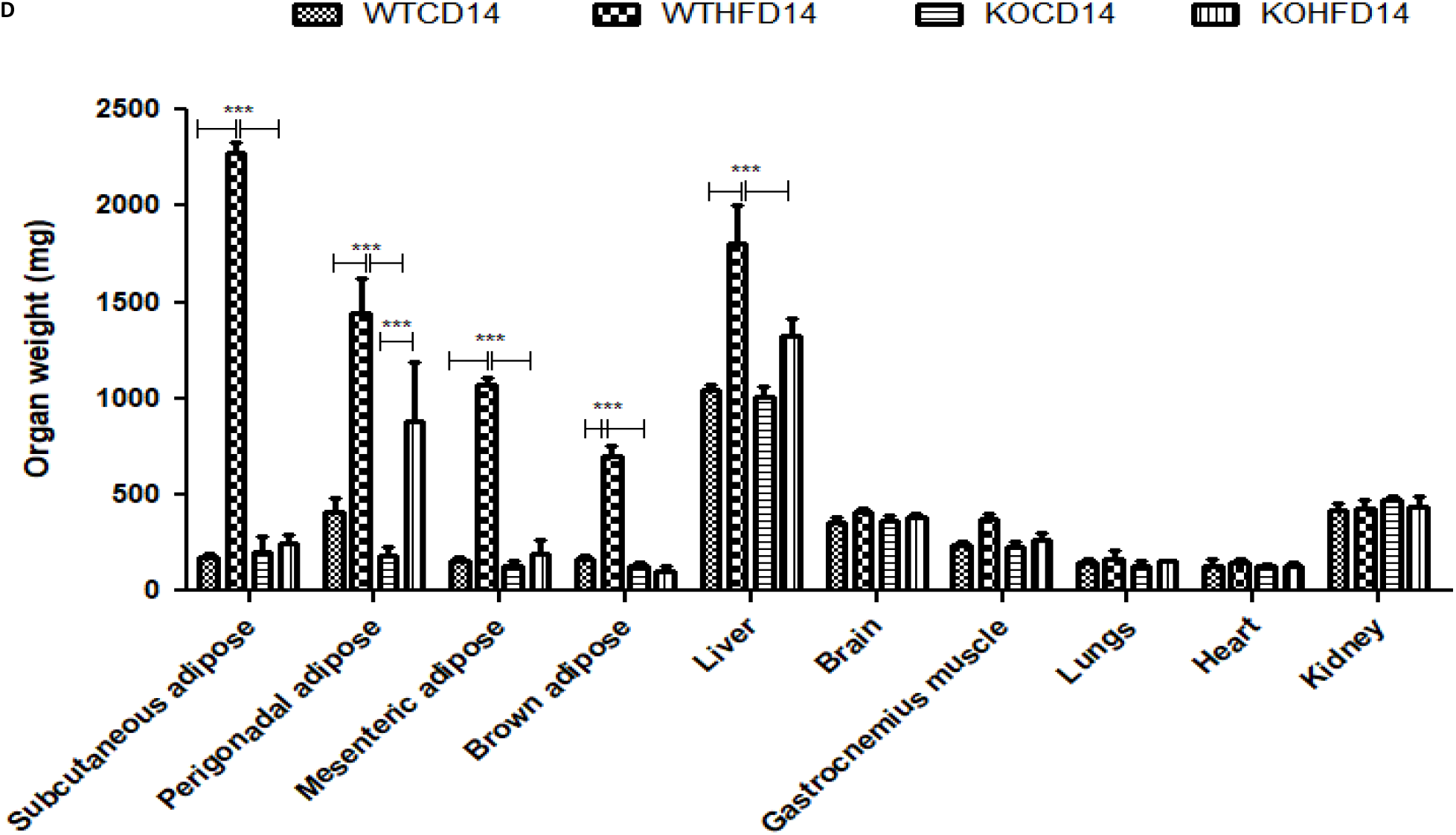
Effect of *slc7a8* deletion on body weight and caloric intake. WTHFD significantly gained weight throughout the 14-week period starting from week 2 when compared to WTCD (p<0.05 to p<0.001). KOHFD significantly gained weight in comparison to KOCD (p<0.05). The difference in weight gain between WTHFD and KOHFD was significant starting week 3, with the p-value increasing gradually from p<0.05 to p<0.001. The WTCD and KOCD showed no differences in weight, A. Total cumulative food consumption was similar across the four groups, except that WTHFD had significantly greater food intake at week 11 when compared to WTCD and KOHFD, B. Energy intake increased significantly from p<0.01 at week 2 to p<0.001 from week 3 to week 14 between WTCD and WTHFD. A significant difference (p<0.001) in caloric intake was observed between WTHFD and KOHFD from week 11 to week 14. No significant differences in caloric intake were seen between KOCD and WTCD, C. WTHFD showed significantly larger (p<0.001) iWAT, pWAT, mWAT, BAT and liver compared to WTCD and KOHFD, D. A-C: Week 1-5: N=18 for WTCD, WTHFD, KOHFD and N=17 for KOCD; Week 6-8: N=12 for WTCD, WTHFD, KOHFD and N=11 for KOCD; Week 9-12: N=6 for WTCD, WTHFD, KOHFD and N=5 for KOCD; Week 13-14: N=5 for WTCD, WTHFD, KOCD and N=6 for KOHFD.

### Deficiency in *slc7a8* had no effect on glucose and insulin metabolism but significantly improved glucose tolerance on HFD

WT and KO mice showed no difference in metabolism of exogenous glucose (Figure 2A and B) and insulin (Figure 2C and D) prior to introducing them on the experimental diets. However, significantly elevated glucose levels (p<0.01) were observed for the KO mice at 30 minutes. After 5 weeks on experimental diets, no significant difference was observed in glucose metabolism between WT and KO on either the CD or HFD (Figure 3A and B). Conversely, at 14 weeks, WTHFD had significantly higher glucose levels compared to KOHFD and WTCD starting from 30 minutes (Figure 3C). Although WTHFD had a larger AUC compared to KOHFD and WTCD, this was not statistically significant (Figure 3D). No significant differences were observed between the AUC of WTCD5, WTHFD5, KOCD5, KOHFD5 when compared to their 14-week counterparts (WTCD14, WTHFD14, KOCD14 and KOHFD14 (Figure 3B and D).

**Figure 2:**
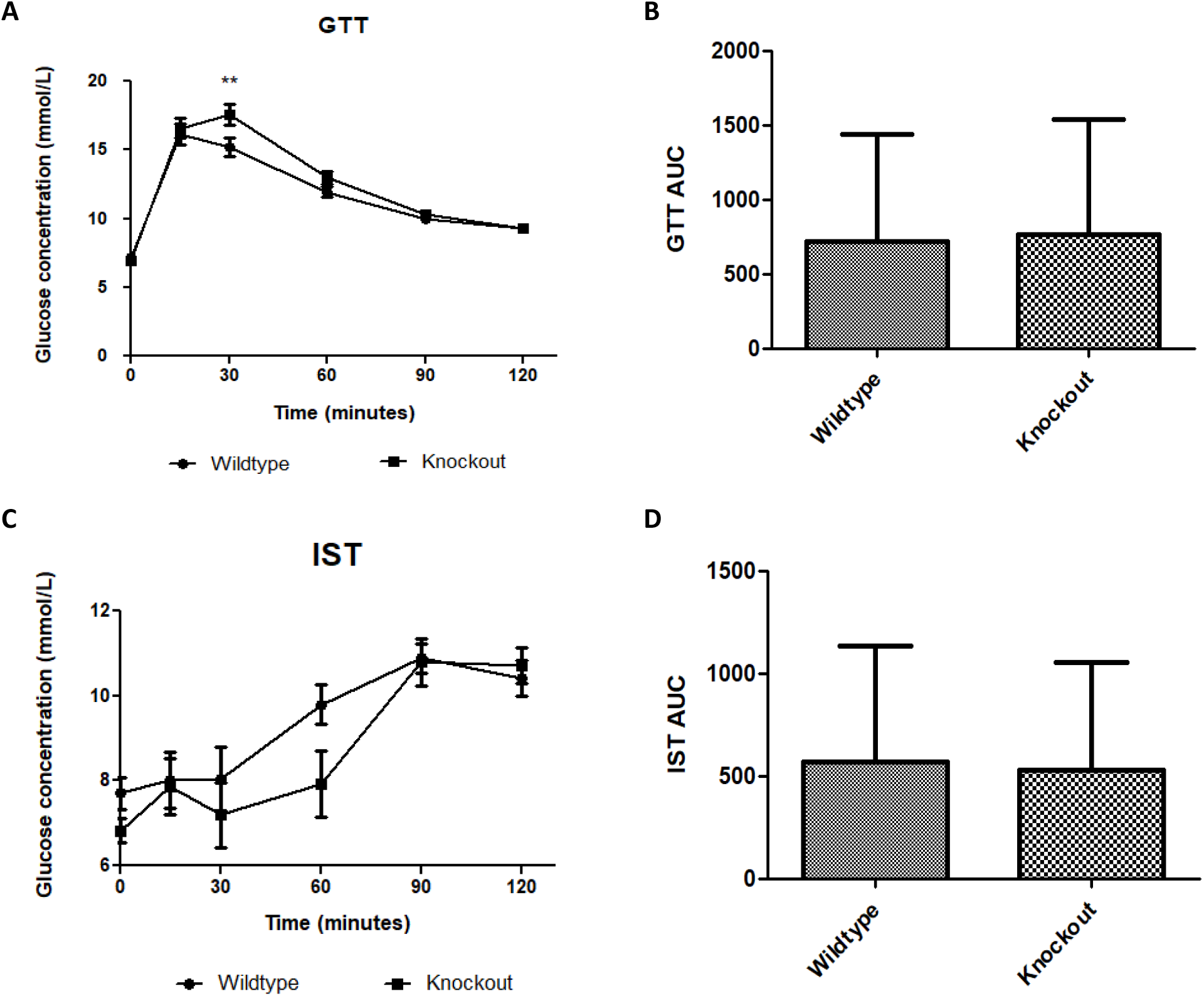
Effect of genotype on glucose tolerance and insulin sensitivity tests. GTT and IST were conducted before introducing the C57BL/6J wildtype and *Slc7a8* knockout mice to CD and HFD. No significant differences were observed in GTT and IST between the WT and KO except that significantly higher glucose levels (p<0.01) were observed for the KO mice at 30 minutes of the GTT, A. GTT: N=47 for WT and N=48 for KO; IST: N=47 for WT and N=44 for KO.

**Figure 3:**
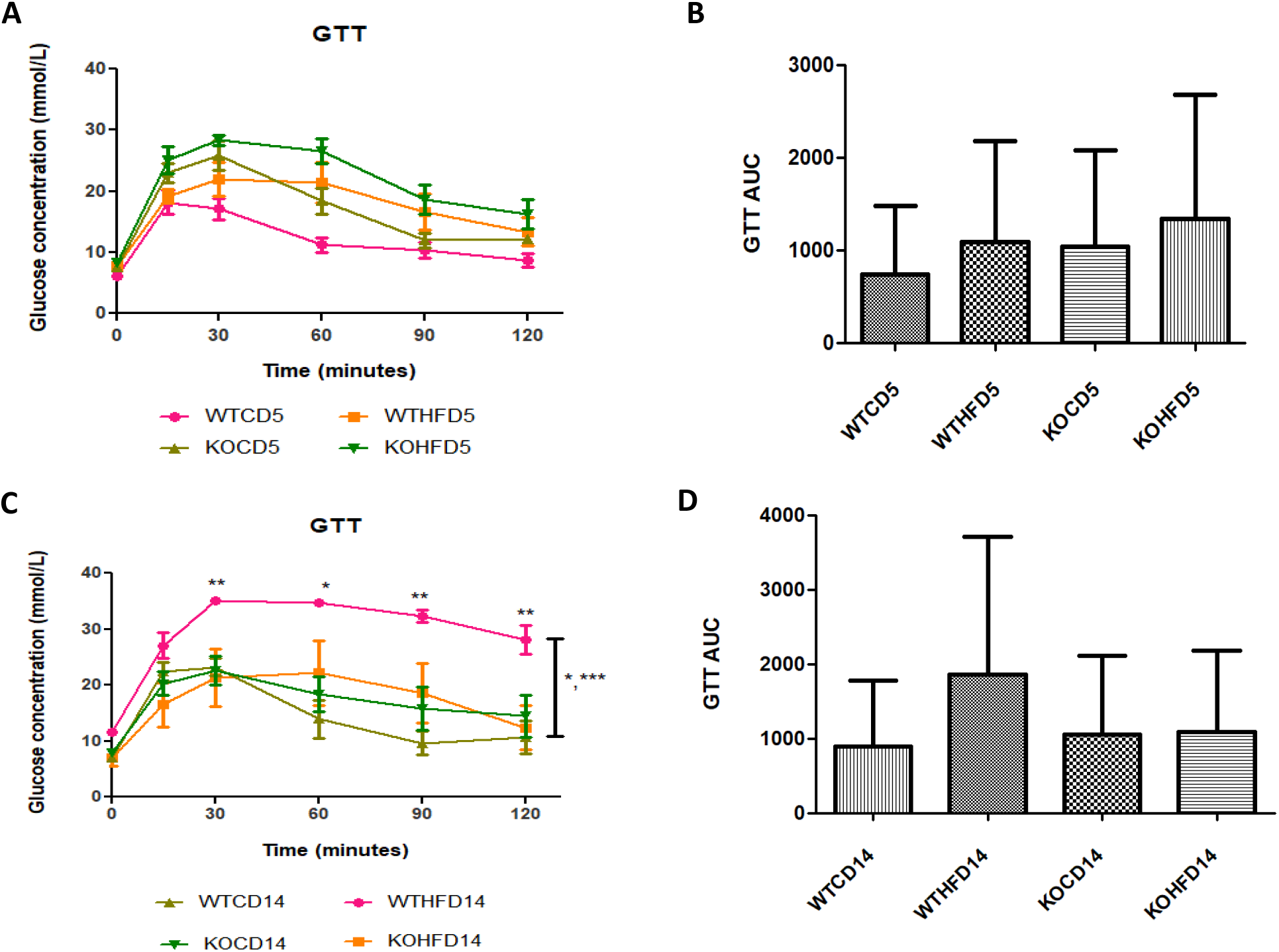
Glucose tolerance and insulin sensitivity tests of animals on experimental diet. No significant differences were observed after experimental feeding between WT and KO on either CD and HFD at 5 weeks, A and B. The WTHFD showed significantly higher glucose levels compared to KOHFD (p<0.05, 0.01) and WTCD (p<0.05, 0.001) at 14 weeks. No significant differences were observed between WTCD5, WTHFD5, KOCD5, KOHFD5 and their respective 14-week counterparts, B and D. N=6 for WTCD5, WTHFD5, KOCD5, KOHFD5, WTCD14 WTHFD14, KOCD14, and N=5 for KOHFD14

### *Slc7a8* deletion attenuates adipocyte hypertrophy in white and brown adipose depots

The pWAT from WTCD (Figure 4A) and KOHFD (Figure 4B) had significantly smaller (p<0.001) adipocyte sizes compared to WTHFD (Figure 4C) as indicated in the column graph (Figure 4D). The number of adipocytes per field was significantly higher in WTCD (p<0.001) and KOHFD (p<0.05) compared to WTHFD (Figure 4E). The iWAT in WTHFD (Figure 4H) had a significant increase (p<0.001) (Figure 4I) in adipocyte hypertrophy compared to KOHFD (Figure 4G) and WTCD (Figure 4F). Similarly, a significant increase (p<0.001) was observed in adipocyte size of mWAT WTHFD (Figure 4M) in comparison to KOHFD (Figure 4L) and WTCD (Figure 4K), Figure 4N. The number of adipocytes per field was significantly lower in mWAT (p<0.01) (Figure 3J) and iWAT (p<0.001) (Figure 3O) of WTHFD compared to WTCD, as well as in mWAT (p<0.01) and iWAT (p<0.001) of WTHFD compared to KOHFD. Lipid droplet accumulation was greater in BAT of WTHFD (Figure 4R) compared to WTCD (Figure 4P) and KOHFD (Figure 4Q). Additionally, as early as 5 weeks on experimental diet, adipocyte hypertrophy was greater in WTHFD compared to KOHFD and WTCD in pWAT, mWAT, iWAT, and larger lipid droplets were observed in BAT of WTHFD (Figure 2 supplementary).

**Figure 4:**
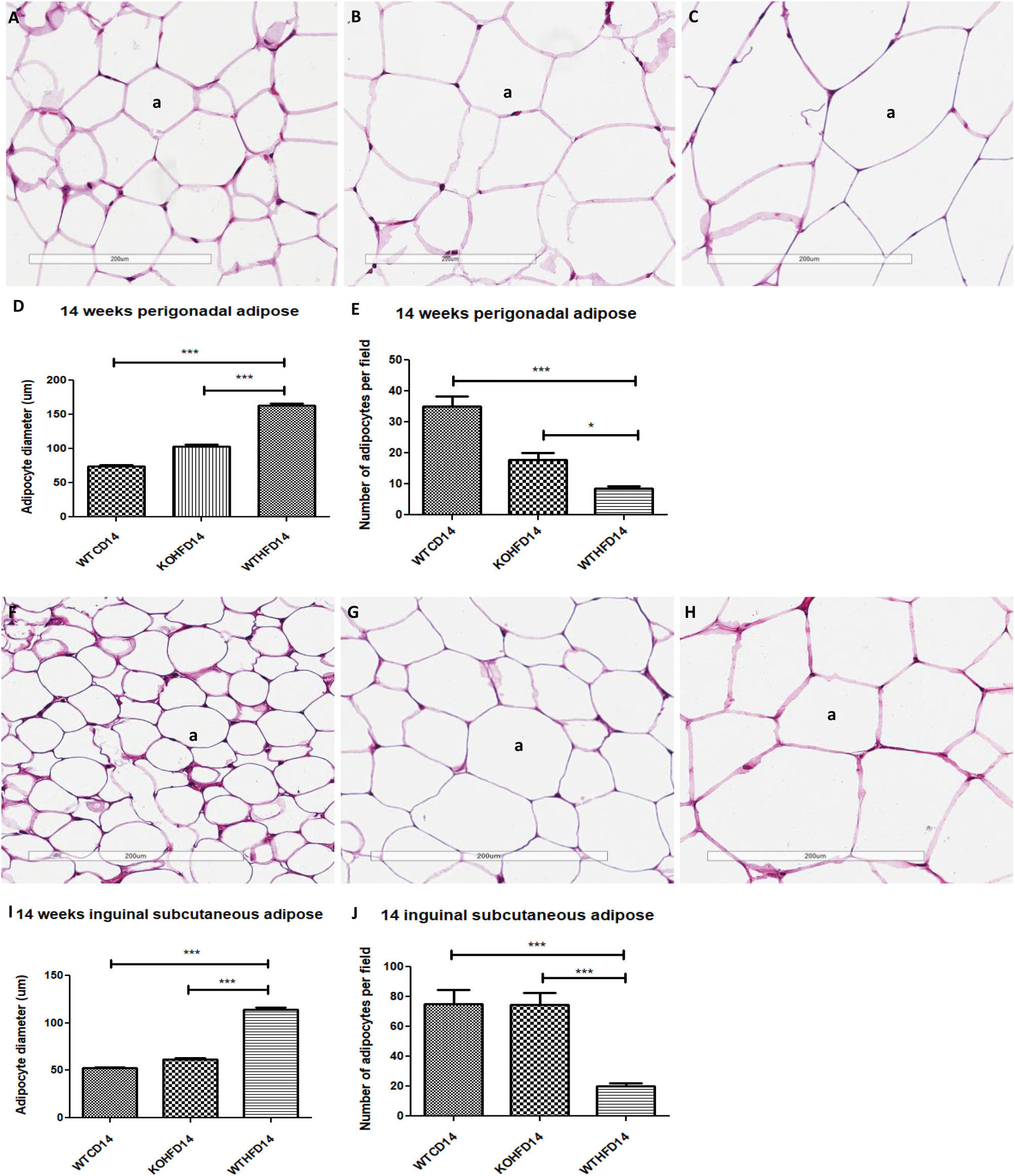
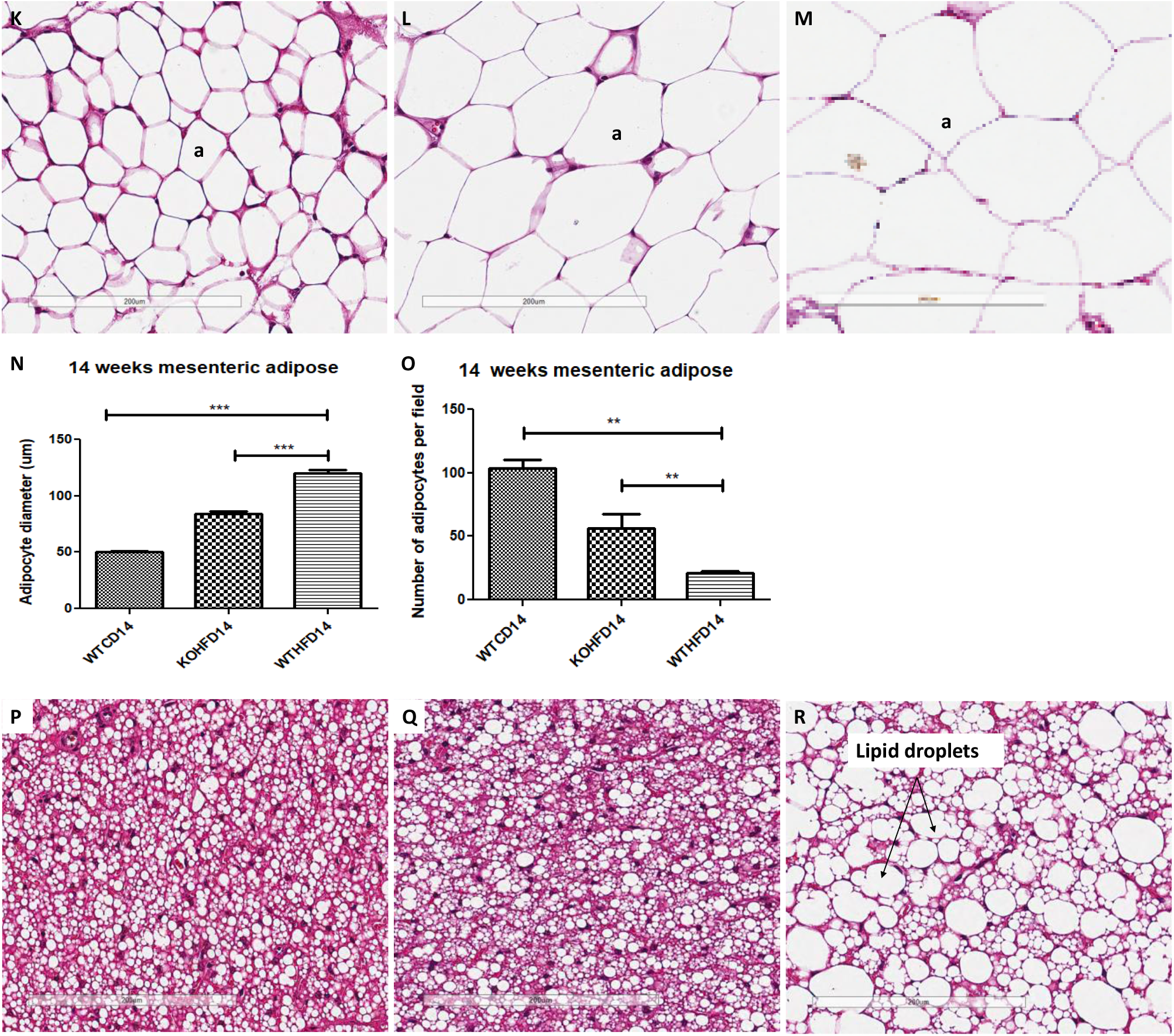
Adipocyte size distribution across the various adipose tissue depots. H&E-stained sections of perigonadal WAT (pWAT) revealed that the WTHFD, C, had significantly larger (p<0.001), D, adipocytes than WTCD, A and KOHFD, B. The number of adipocytes per field was significantly smaller in WTHFD than KOHFD (p<0.05) and WTCD (p<0.001), E. Similarly, adipocyte diameter of WTHFD, H, of inguinal subcutaneous WAT (iWAT) was significantly greater (p<0.001), I, than that of WTCD, F, and KOHFD, G. Conversely, the number of adipocytes per view was significantly lower (p<0.001) in WTHFD compared to WTCD and KOHFD, J. Significant (p<0.001), N, adipocyte hypertrophy was also observed in mWAT of WTHFD, M, compared to WTCD, K and KOHFD, L. Additionally, significantly (p<0.01) fewer adipocytes were viewed per field in WTHFD compared to WTCD and KOHFD, O. Sections of the BAT revealed that WTCD, P, and KOHFD, Q, had smaller lipid droplets compared to those observed in WTHFD, R. Magnification= 20X, Scale bar= 200 μm. Key: a= adipocyte. N =120 adipocytes

### Deletion of *slc7a8* reduces liver steatosis in diet induced obese mice

Liver sections from WTHFD (Figure 5C) showed lipid accumulation which can be categorised as microvesicular (circled, Figure 5C) and macrovesicular (indicated in arrow, Figure 5C) steatosis. This phenomenon was absent in liver sections of WTCD (Figure 5A). While macrovesicular steatosis was observed in KOHFD (Figure 5B), the lipid droplets were visibly smaller when compared to those observed in WTHFD. Congestion of the central vein was observed in WTHFD (Figure 5F) but not in WTCD (Figure 5D) and KOHFD (Figure 5E). No visible changes in sinusoid dilation and Kupffer cell morphology were observed between WTCD, KOHFD and WTHFD. Additionally, lipid droplets in the form of micro- and macrovesicular steatosis were observed as early as 5 weeks in WTHFD, while macrovesicular steatosis was also seen in the KOHFD but was visibly smaller comparison to WTHFD (Figure 3 supplementary).

**Figure 5:**
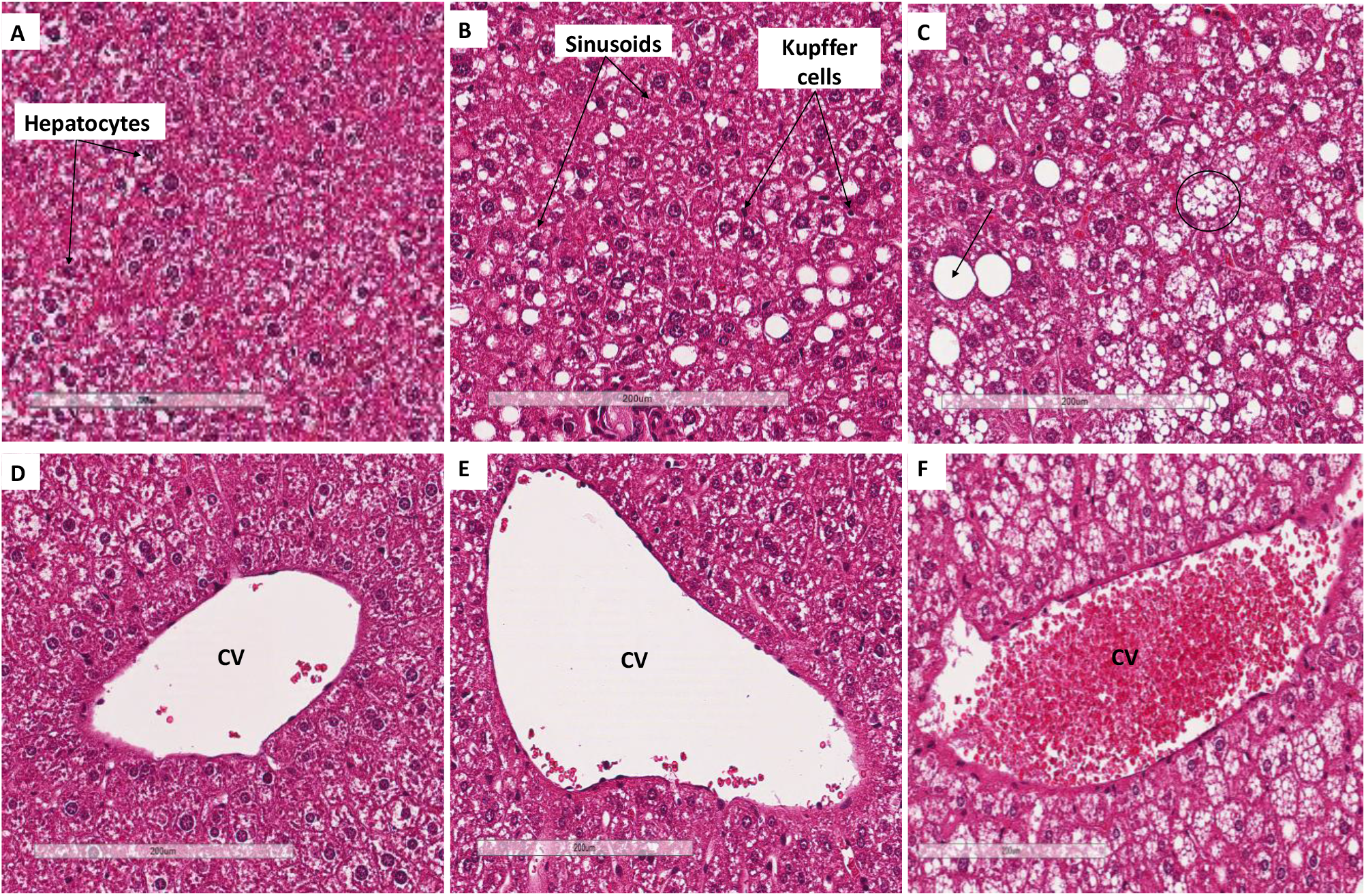
Lipid accumulation in the liver. H&E-stained liver sections showed the presence of micro- and macrovesicular steatosis in WTHFD, C which was not observed in WTCD, A and KOHFD, B. Venous congestion in WTHFD, F was also observed but not in WTCD, D and KOHFD, E. Magnification = 20X, Scale bar= 200 μm. Key: CV=central vein

### Deficiency in *slc7a8* decrease lipid accumulation in gastrocnemius muscle

Myocyte atrophy was observed in WTHFD (Figure 6C) which had significantly smaller myocytes (p<0.001) (Figure 6G) than WTCD (Figure 6A). The deletion of *slc7a8* increases myocyte size in KOHFD (Figure 6B) compared to WTHFD (Figure 6G). Accumulation of peri-muscular adipose tissue (PMAT) (Figure 6F) was observed to be greater in WTHFD than in KOHFD (Figure 6E) and WTCD (Figure 6D). At week 5, the KOHFD had significantly larger myocytes (p<0.001) and less adipose tissue accumulation than WTHFD (Figure 4 supplemetary).

**Figure 6:**
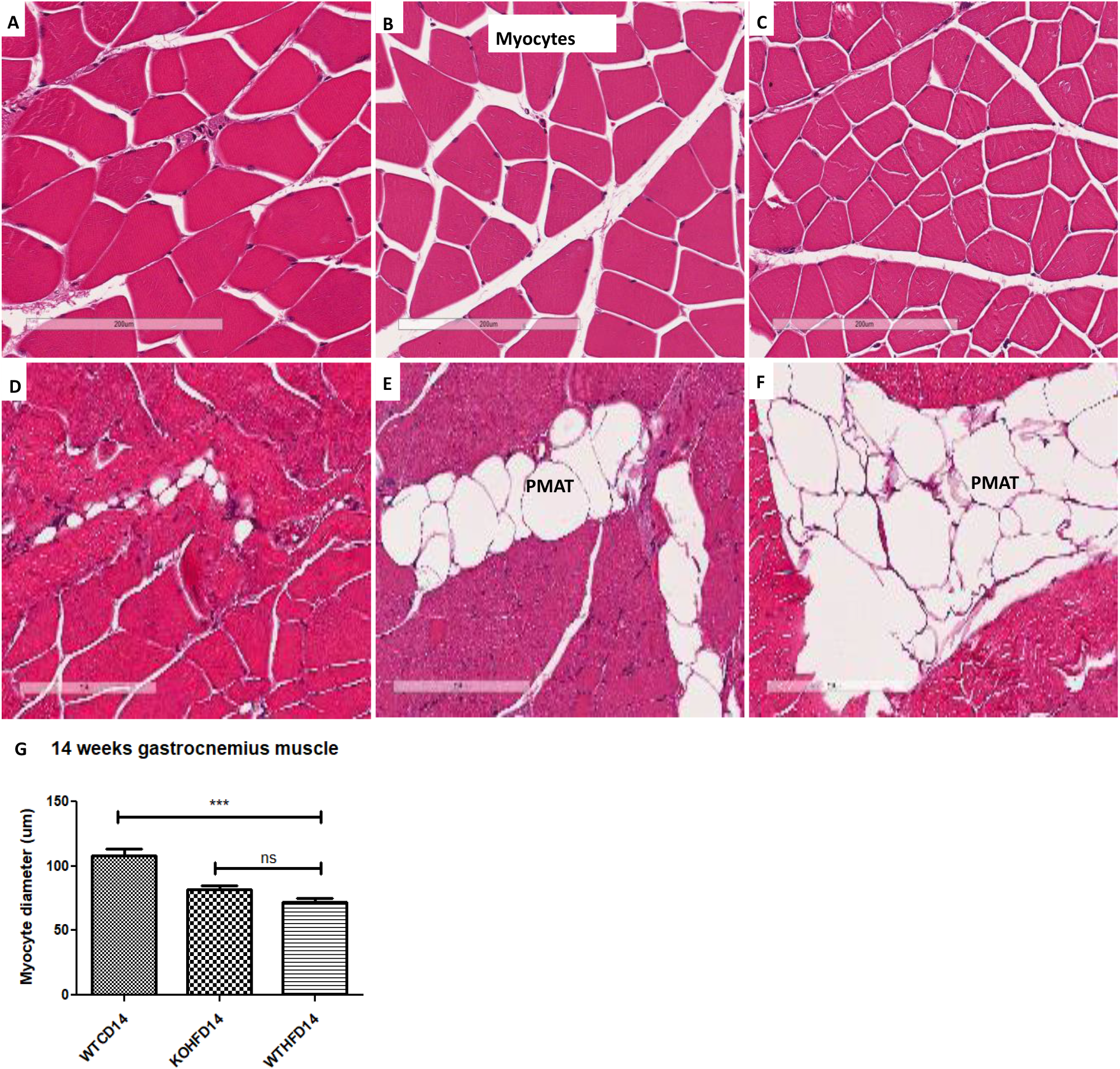
Effect of *slc7a8* deletion on adipose tissue accumulation and myocyte atrophy in gastrocnemius muscle. KOHFD (Figure 6B) resulted in a protective effect against muscle atrophy when compared to WTHFD (Figure 6C). WTHFD had significantly smaller (p<0.001) (Figure 6G) myocytes than WTCD (Figure 6A). Greater peri-muscular adipose tissue (PMAT) accumulation was seen in WTHFD (Figure 6F) when compared to WTCD (Figure 6D) and KOHFD (Figure 6E). N= 120 myocytes

### Deficiency in *slc7a8* reduces accumulation of epicardial adipose tissue

The increase in the accumulation of epicardial adipose tissue (EAT - white adipose tissue) observed in WTHFD (Figure 7C) compared to WTCD (Figure 7A) was decreased following the deletion of *slc7a8*, KOHFD (Figure 7B). Larger lipid droplets were observed in brown/beige adipose tissue (a property of epicardial adipose tissue) in WTHFD (Figure 7F) when compared to WTCD (Figure 7D) and KOHFD (Figure 7E). The connective tissue of the WTHFD (Figure 7I) was visibly thicker than that of the WTCD (Figure 7G) and KOHFD (Figure 7H). Additionally, greater accumulation of adipose tissue (black arrow) was observed surrounding the coronary artery (CA) of WTHFD (Figure 7L) compared to KOHFD (Figure 7K) and WTCD (Figure 7J).

**Figure 7:**
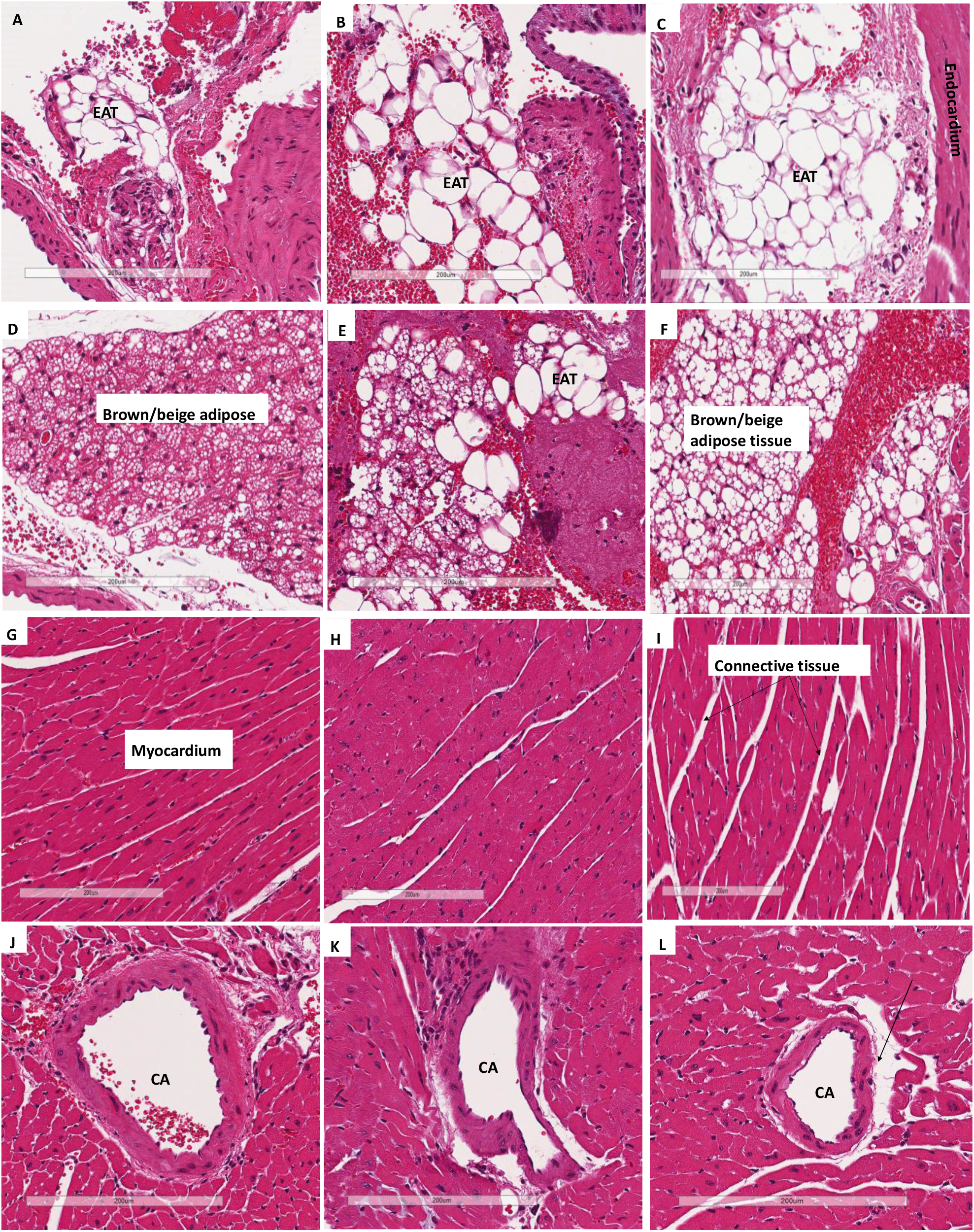
Effect of *slc7a8* on epicardial adipose tissue accumulation in the heart. H&E-stained heart sections showed a greater accumulation of epicardial adipose tissue, seen as a brown/beige adipose depot, in the WTHFD, C, compared to WTCD, A and KOHFD, B. The images demonstrate that the WTHFD, F, mice had more connective tissue (cardiac muscle fibres) than WTCD, D and KOHFD, E. Additionally, the coronary artery of the WTHFD, I, was surrounded by larger lipid deposits (black arrow) in comparison to WTCD, G and KOHFD, H. Magnification = 20X, Scale bar= 200 μm. Key: CA=coronary artery

### Deficiency in *slc7a8* reduces lipid accumulation in the ganglion layer in diet induced obesity

Brain tissues of WTHFD (Figure 8C) showed vacuolation in the Purkinje cell layer (indicated by black arrows) when compared to WTCD (Figure 8A). Deletion of *slc7a8* attenuated the vacuolation observed in DIO, KOHFD (Figure 8B). In the cerebral cortex, lipid droplets were seen in KOHFD (Figure 8E) and WTHFD (Figure 8F), but not in WTCD (Figure 8D). No visible morphological differences were observed between Purkinje cells in WTCD (Figure 8A), KOHFD (Figure 8B) and WTHFD (Figure 8C).

**Figure 8:**
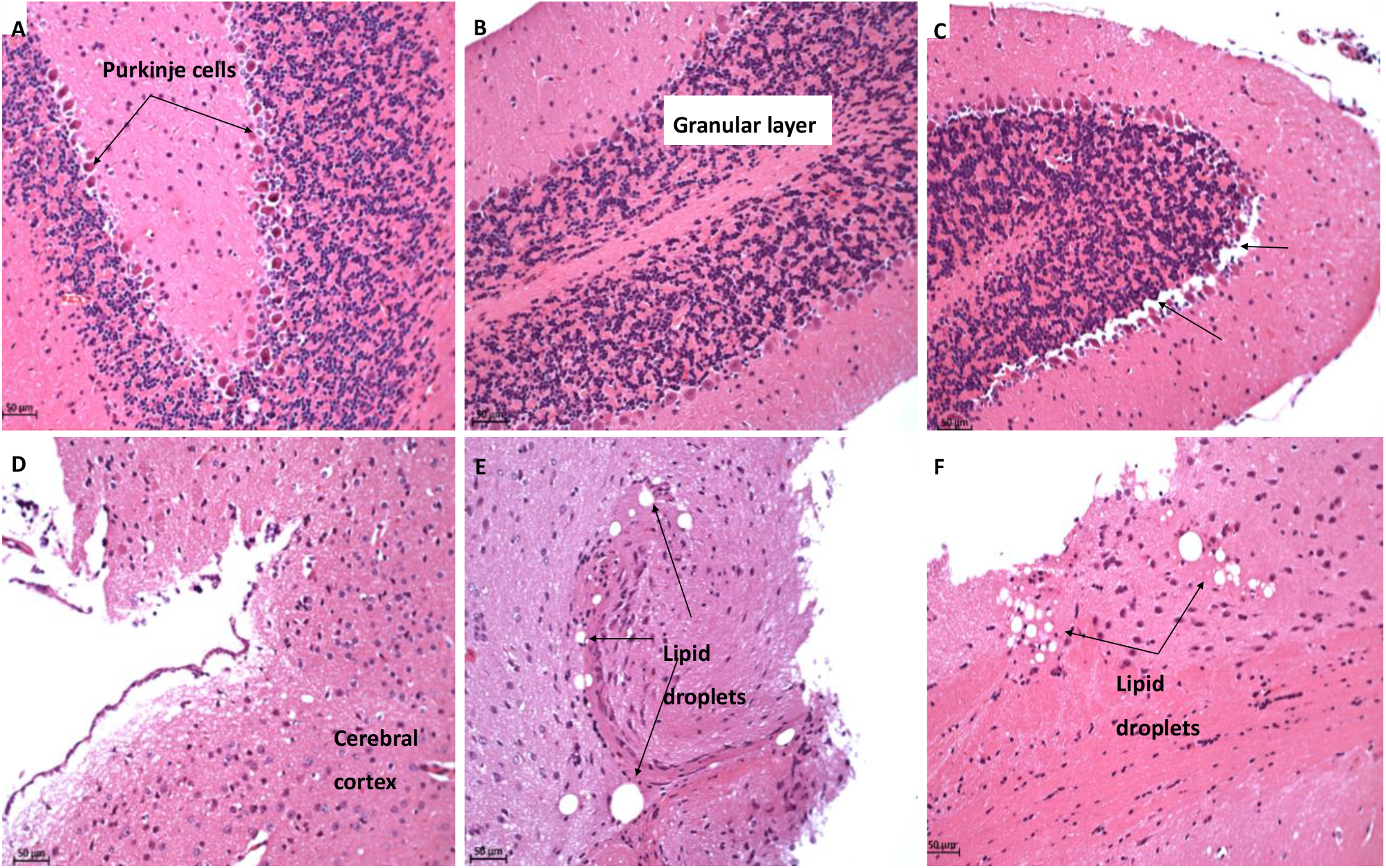
Effect of *slc7a8* deletion on lipid droplet accumulation in brain tissue. H&E-stained sections of brain tissue showed vacuolation in the Purkinje cell layer of WTHFD, C, when compared to WTCD, A and KOHFD, B. Lipid droplets were observed in the cerebral cortex of KOHFD, E and WTHFD, F, which was not seen in WTCD, D. Magnification = 20X, Scale bar= 50 μm

### Deficiency in *Slc7a8* reduces glomerulus size and lipid accumulation in the kidney

No visible alterations were seen in the renal tubules of WTCD (Figure 9A), KOHFD (Figure 9B) and WTHFD (Figure 9C). However, WTHFD had enlarged glomeruli which were significantly larger (p<0.01) than those of WTCD. Although glomerular sizes in KOHFD (Figure 9B) appeared to be smaller than those of WTHFD, the difference was not statistically significant (Figure 9G). The Bowman’s space of WTHFD (Figure 9F) was significantly larger (p<0.05) with the presence of many larger lipid droplets than in WTCD (Figure 9D). Interestingly, the deletion of *slc7a8* significantly reduced the Bowman’s space enlargement and lipid droplet accumulation in DIO, KOHFD (Figure 9H) to a level observed in WTCD (Figure 9 D and H).

**Figure 9:**
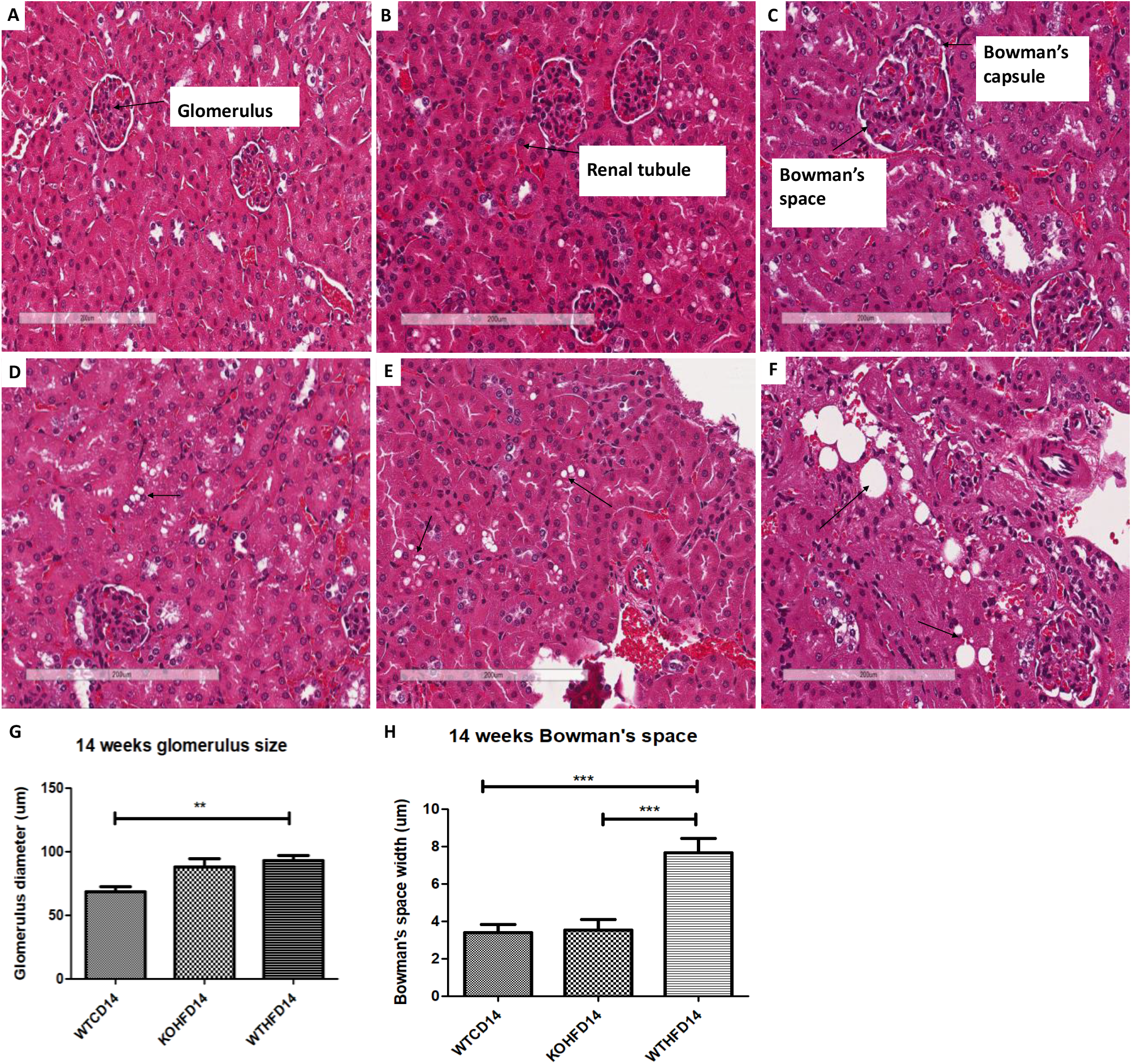
Effect of *slc7a8* deletion on lipid accumulation and glomerular size in the kidneys. H&E-stained sections showed that WTHFD, C, had significantly enlarged glomeruli (p<0.01), G, compared to WTCD, A; no significant differences were observed between glomeruli of WTHFD and KOHFD, C. The width of the Bowman’s space was significantly larger (p<0.05) in WTHFD when compared to WTCD and KOHFD, H. Accumulation of lipid (black arrows) was greater in WTHFD, F, when compared to WTCD, D and KOHFD, F. Magnification = 20X, Scale bar= 200 μm. N= 10 for glomeruli and bowman’s space

### Deficiency in *slc7a8* reduces adipose tissue accumulation in the lungs

Histological analysis of the lungs showed that adipose tissue tends to lie adjacent to bronchioles and pulmonary arteries in WTCD (Figure 10A and D), KOHFD (Figure 10B) and WTHFD (Figure 10C). In DIO, greater accumulation of adipose tissue was observed in WTHFD (Figure 10C) when compared to WTCD (Figure 10A). The accumulation of adipose tissue in DIO appeared to reduce in KOHFD (Figure 10B). The smooth muscle layer was enlarged in WTHFD (Figure 10F) in comparison to WTCD (Figure 10D) and KOHFD (Figure 10E). Additionally, the alveolar walls around the alveolar sacs (AS) of the WTHFD (Figure 10I) appeared thinner in comparison to WTCD (Figure 10G). Interestingly, the deletion of *slc7a8* abated this effect that resulted from DIO, in KOHFD (Figure 10H). Lipid accumulation in the lungs was observed as early as week 5 with more adipose tissue in WTHFD and KOHFD compared to WTCD (Figure 5 supplementary).

**Figure 10:**
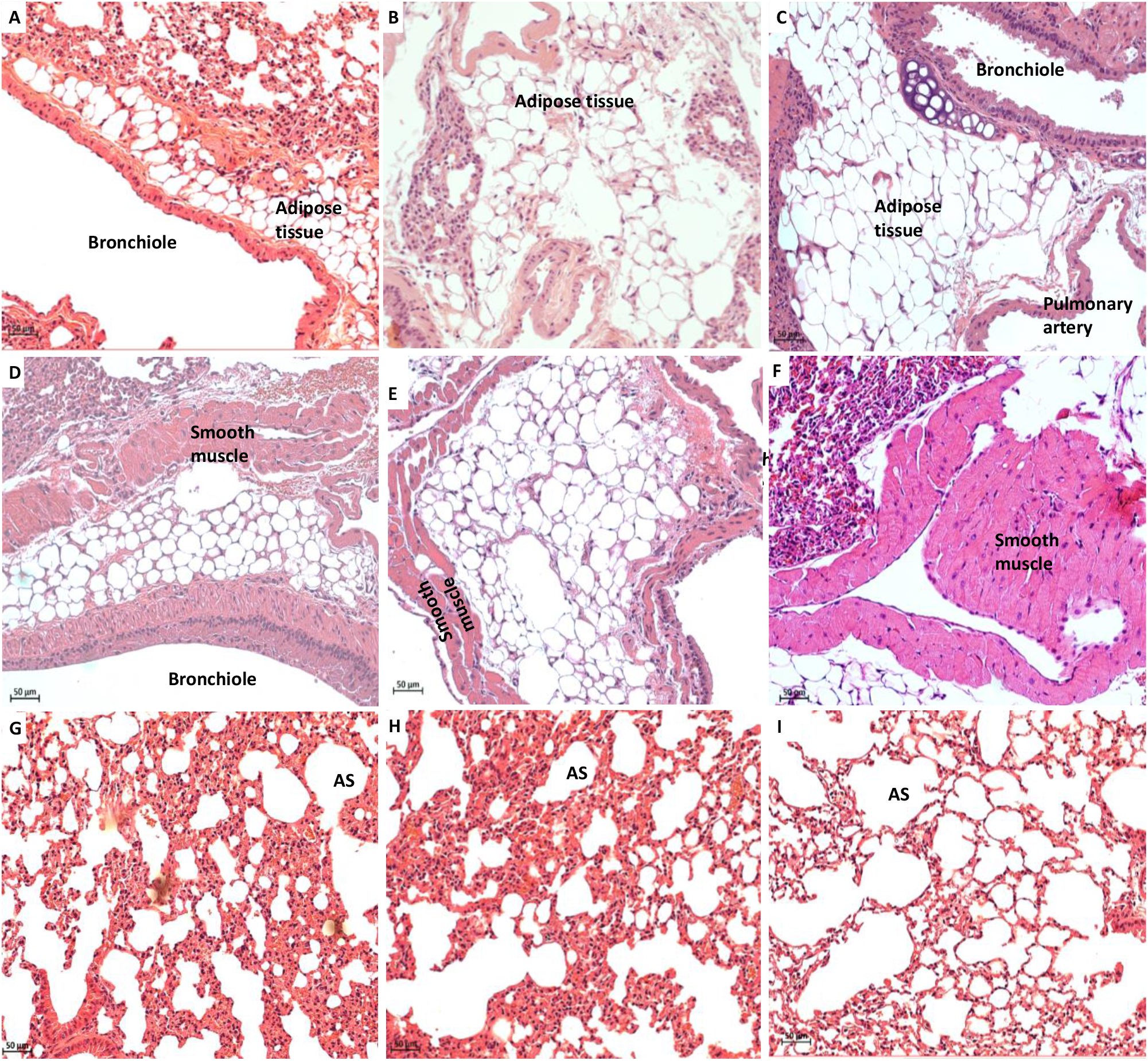
Effect of *slc7a8* deletion on lipid accumulation in the lungs. H&E-stained lung sections showed the accumulation of adipose tissue around the bronchioles and pulmonary artery, which was greater in WTHFD, C, compared to KOHFD, B, and WTCD14, A, D. An enlarged smooth muscle layer was observed in WTHFD, F compared to KOHFD, E and WTCD, D. Additionally, thinner alveolar walls were observed in WTHFD, I, in comparison to WTCD, G, and KOHFD, H. Magnification = 20X, Scale bar= 50 μm. Key: AS= Alveolar sacs

### Deficiency in s*lc7a8* reduces white adipose tissue inflammation in DIO

Immunohistochemical staining for F4/80, a mouse macrophage marker, was done to assess the presence of macrophages in pWAT, mWAT, iWAT and brown adipose tissue. Deletion of *slc7a8* significantly decreased macrophage infiltration (indicated by black arrows) in the pWAT (Figure 11C; p<0.01), mWAT (Figure 11F; p<0.05) and iWAT (Figure 11I; p<0.01) of KOHFD (Figure 11A, D & G) compared to WTHFD (Figure 11B, E & H). No significant difference was observed in the brown adipose macrophage inflammation profile (Figure 11L) between WTHFD (Figure 11K) and KOHFD (Figure 11J).

**Figure 11:**
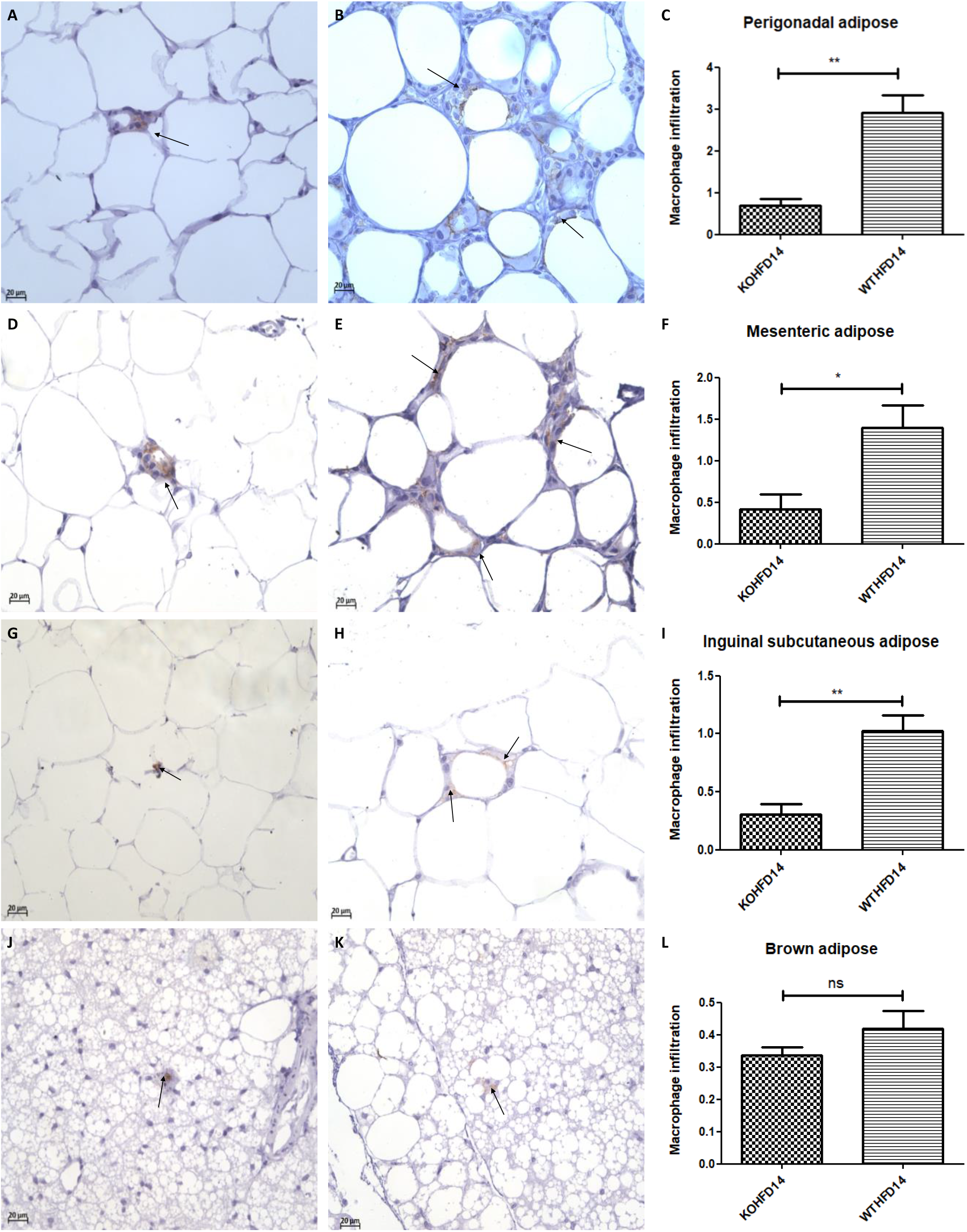
Effect of *slc7a8* deletion on macrophage infiltration in adipose tissues. KOHFD, A, showed a significant decrease in macrophage infiltration (indicated by black arrows) in the pWAT (p<0.01), C, compared to WTHFD, B. A significant decrease in macrophage infiltration in mWAT (p<0.05), F, was observed in KOHFD, D, compared to WTHFD, E. A significantly decrease in macrophage infiltration (p<0.01), I, in iWAT was seen in KOHFD, G, when compared to WTHFD, H. No significant differences in macrophage infiltration in brown adipose tissue, L, was observed between the KOHFD, J and WTHFD, K. Magnification = 40X, Scale bar= 20 μm. N= 5 fields

### *Deficiency in slc7a8* reduces inflammation in the liver

Deficiency in *slc7a8* resulted in a significant (p<0.05) reduction in macrophages in the liver in DIO (KOHFD), Figure 12A, compared to WTHFD (Figure 12B), Figure 12C.

**Figure 12:**
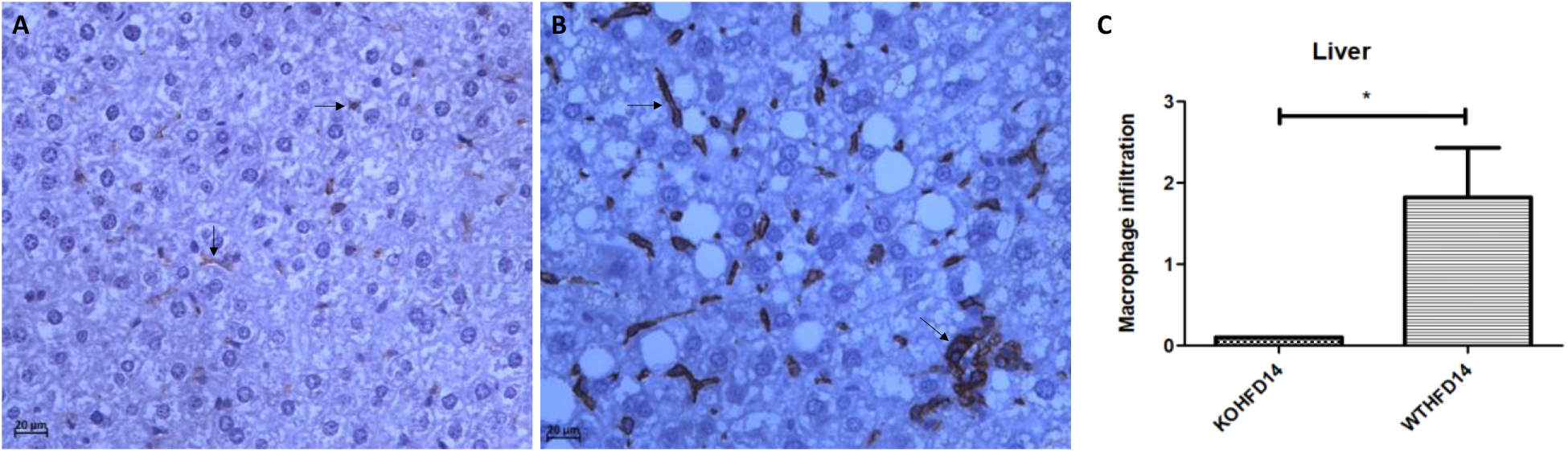
Effect of slc7a8 deletion on the presence of macrophages in the liver. WTHFD, B, had a significantly greater infiltration (p<0.05) of macrophages compared to KOHFD, A, C. Magnification = 40X, Scale bar= 20 μm. N= 10 fields

### Deficiency in *slc7a8* had no effect on the presence of macrophages in kidney and gastrocnemius muscle in DIO

The presence of macrophages in the kidney of KOHFD (Figure 13A) and WTHFD (Figure 13B) was similar (Figure 13C). This observation was the same for gastrocnemius muscle of KOHFD (Figure 13D) and WTHFD (Figure 13E) with no statistical difference in macrophage profile between them (Figure 13F).

**Figure 13:**
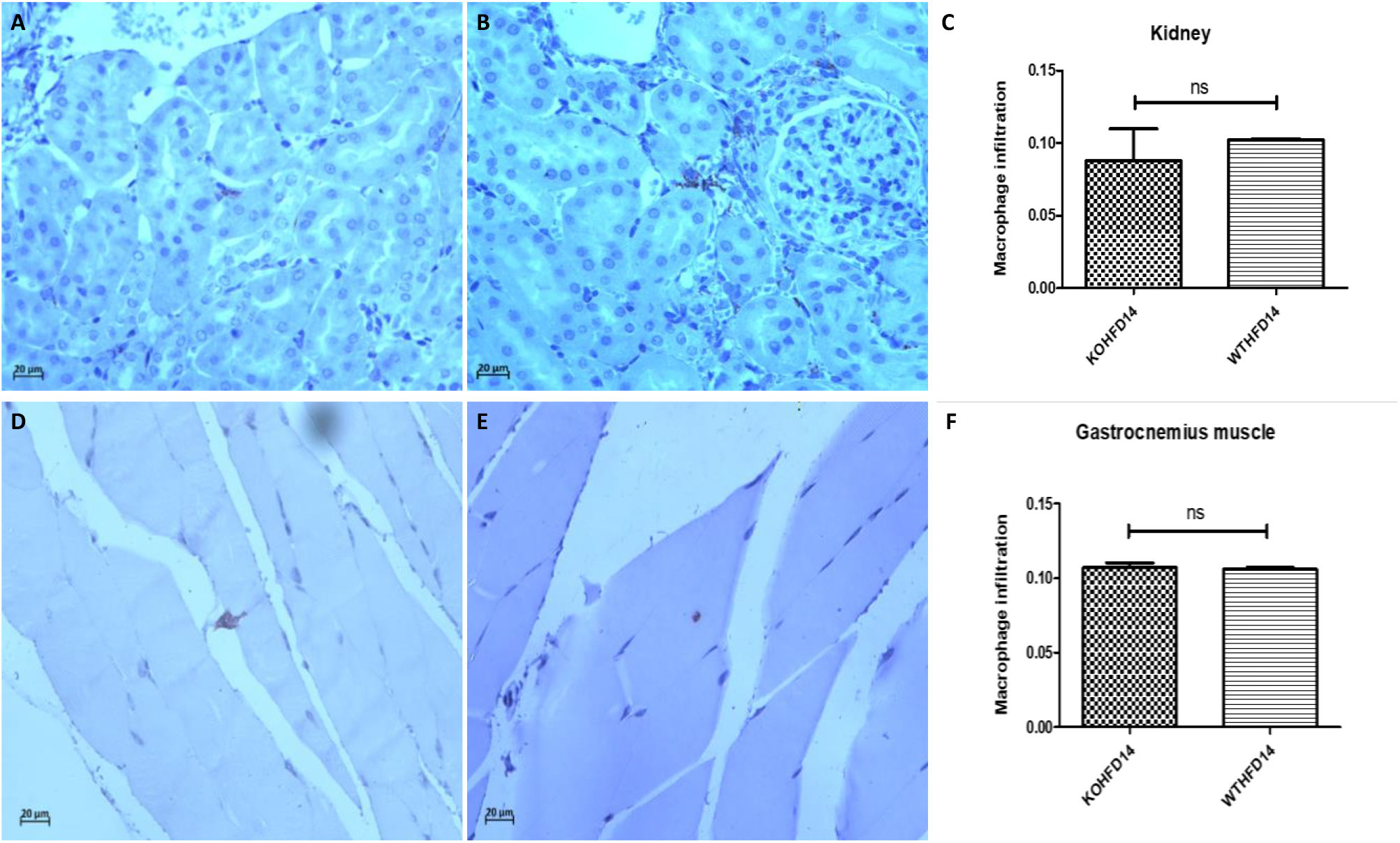
Effect of *slc7a8* on macrophage infiltration profile of kidney and gastrocnemius muscle. KOHFD, A, had slightly fewer macrophages infiltrating into the kidney in comparison to WTHFD, C. However, no significant differences were noted between KOHFD and WTHFD. No significant differences, F, were observed in infiltration between the gastrocnemius muscles of KOHFD, D and WTHFD, E. Magnification = 40X, Scale bar= 20 μm. N= 5 fields

## Discussion

Obesity is characterized by excessive accumulation of adipose tissue, and it is associated with the development of metabolic syndromes affecting many organs and tissues in the body. The search for molecular factors that play a role in attenuating lipid accumulation in condition such as diet induced obesity is paramount to identifying good candidates for therapeutic interventions that mitigate the development of obesity associated comorbidities. Studies of adipogenesis in human derived stromal/stem cells *in vitro* have served as an excellent model for identifying molecular factors with a potential role in adipocyte formation and lipid accumulation/metabolism [11, 14]. This study investigated the role of a previously identified novel human adipogenic gene, SLC7A8 [14]in diet-induced obesity, and its effect on adipose tissue accumulation in different organs and tissues. To achieve this, *slc7a8* knockout (KO) and wildtype (WT) C57BL/6 mice were fed either a HFD or nutrient matched CD for 14 weeks followed by the analyses of different parameters.

Weight gain, food, and caloric intake between WTCD and KOCD were similar, indicating that *slc7a8* deletion had no effect on food intake, caloric consumption, and weight gain on a normal diet. WTHFD gained significantly more weight (p<0.001) than WTCD starting from week 3 (Figure 1A) with a significantly higher caloric intake (p<0.01 to p<0.001) than WTCD (Figure 1C). Total food consumption was not significantly different during the 14-week period except at week 8 where food consumption in WTHFD was significantly elevated (p<0.05). This indicates that the occurrence of diet-induced obesity was due to an increase in caloric intake when on HFD. Interestingly, the *slc7a8* deficient genotype on HFD (KOHFD) gained significantly less weight (p<0.05 to p<0.001) compared to the WTHFD starting from week 3 (Figure 1A). This suggests that *slc7a8* deletion is protective against diet-induced obesity. The significant decrease in weight gain in KOHFD was accompanied by significantly lower tissue mass of iWAT, mWAT, pWAT, BAT and liver compared to WTHFD (Figure 1D). Strikingly, it was observed that KOHFD gained significantly more weight (p<0.05 to p<0.001) than KOCD from week 8, and this corresponded to a significantly larger pWAT in KOHFD than KOCD (Figure 1D). This indicates that weight gain by KOHFD is due to pWAT expansion and suggests that pWAT is the primary site of lipid accumulation in the KO phenotype.

BAT in WTHFD (Figure 4R) displayed enlarged lipid droplets compared to WTCD (Figure 4P). A recent study showed that following 20 weeks of feeding mice on a HFD, lipid accumulation did not influence the function of brown adipose tissue. However, the authors speculated that if the period of HFD feeding was extended, a malfunction of BAT would be observed in obese mice[16]. We have observed in this current study that KOHFD (Figure 4Q) attenuates adipocyte hypertrophy and lipid accumulation in BAT. This suggests that *slc7a8* deletion could be protective against the long-term effect of BAT hypertrophy and malfunctioning caused by DIO.

Furthermore, it was observed that WTHFD had a significantly greater caloric intake than KOHFD (Figure 1C) while food consumption was similar except at week 11 where a significant difference (p<0.05) was observed. It is possible that the deletion of *slc7a8* regulates weight gain on HFD by burning calories quicker than WTHFD since both KOHFD and WTHFD had similar caloric intake up to week 8 (Figure 1C) but as early as week 5, adipocyte hypertrophy was already significantly greater in WTHFD compared to KOHFD (Figure S2). Additionally, food and caloric intake was similar between KO and WT on a normal diet, with differences only being observed on HFD; this could suggest satiety in KOHFD as caloric intake significantly decreased after week 8 (Figure 1C).

Adipose tissue expansion in obesity is commonly associated with conditions such as hyperglycaemia, impaired glucose tolerance and insulin resistance[17]. To investigate the effect of *slc7a8* deletion on the metabolism of exogenous glucose and insulin, GTT and IST were performed on all animals (KO and WT) prior to introducing them to an experimental diet (Figure 2A and 2B). Importantly, there was no significant difference between the *slc7a8* KO and WT mice for both tests. This shows that the deletion of *slc7a8* had no effect on their ability to metabolise glucose and insulin efficiently. It was noted, however, that significantly higher levels of blood glucose were seen in KO mice at 30 minutes of the GTT (Figure 2A), which later return to normal without any change in the AUC between KO and WT (Figure 2B). Both WTCD and KOCD at 5 and 14 weeks showed a similar trend in glucose metabolism (Figure 3A and C) with no difference in the AUC (Figure 2B and D), suggesting glucose metabolism is unaltered in *slc7a8* deficient mice on a normal diet. Under condition of DIO, WTHFD showed significantly higher levels of glucose intolerance compared to WTCD, and this effect was significantly improved in KOHFD with blood glucose levels returning to baseline levels at the end of the GTT (Figure 3C). This demonstrates that *slc7a8* deletion significantly improves glucose metabolism in DIO.

WTHFD showed significantly larger adipocytes in the pWAT, mWAT and iWAT (Figure 4) compared to WTCD. The adipose tissue hypertrophy in WTHFD may increase susceptibility to hyperglycaemia. In an obese phenotype, insulin signalling is usually impaired, which results in reduced glucose uptake by muscles and thus increased glucose levels in the circulation[18]. pWAT is significantly larger (p<0.001) than iWAT and mWAT in WTHFD (Figure 6 supplementary), which may be suggestive of pWAT being the main site of lipid accumulation in this group as was observed in the KO group. Abdominal/visceral obesity is critical to the development of metabolic syndrome, and accumulation of adipose tissue in the abdomen correlates with metabolic syndrome, compared to lipid accumulation in the subcutaneous depot[19]. Larger pWAT in WTHFD may thus be responsible for the glucose intolerance observed in these mice. Lipid accumulation in the liver presented as microvesicular steatosis (characterised by small lipid droplets in the cytoplasm of hepatocytes) and macrovesicular steatosis (large lipid droplets) (Figure 5), are both of which important in the development of non-alcoholic fatty liver disease (NAFLD)[20, 21], and were observed in WTHFD but not in WTCD. The presence of lipid droplets in WTHFD liver could be due to the redistribution of excess lipid to peripheral organs such as the liver or muscles seen in the obese phenotype, when the storage capacity of adipose tissue is exceeded[3, 5]. The liver has previously been reported to be the major site for storage of free-fatty acids (FFA) released from white adipose tissue in an obese phenotype[22]. Furthermore, the vast majority of hepatic triglycerides in obese individuals with NAFLD are from FFA released from adipose tissue[23]. The observations made in our study indicate that KOHFD attenuates both macrovesicular and microvascular steatosis seen in WTHFD, suggesting that *slc7a8* deletion could be protective against NAFLD in DIO.

DIO is often associated with the recruitment and accumulation of macrophages in adipose depots. The F4/80 antibody is a marker for macrophages in mouse tissues[10, 24] and was utilised in this study. Adipose tissues from obese WTHFD mice showed significantly more macrophages, which indicates increased inflammation when compared to KOHFD (Figure 11). Thus, *slc7a8* deletion significantly improves the inflammatory profile of adipose tissues in DIO. The liver tissue sections in WTHFD showed significantly elevated levels of macrophages and congestion of the central vein. The observed histopathological changes in liver which occur due to DIO were improved by *slc7a8* deletion in KOHFD (Figure 12).

Apart from metabolic syndromes that are associated with excess adipose tissue accumulation, obese individuals are also prone to developing pulmonary disorders such as chronic obstructive pulmonary disease (COPD) or asthma[25]. In DIO, the lungs of WTHFD showed an increase in adipose tissues accumulation around the bronchioles and pulmonary arteries, which was reduced in KOHFD (Figure 10). Additionally, the smooth muscle layer was visibly thicker, and alveolar walls thinner in the WTHFD in comparison to KOHFD (Figure 10). A previous study showed that accumulation of adipose tissue in the lungs increased with an individual’s body mass index (BMI)[26]. Additionally, an increase in adipose tissue affects the structure of the lungs, resulting in the blockage of airways and causing inflammation which ultimately gives rise to pulmonary disease[25, 26]. We observed that the deletion of *slc7a8* attenuates adipose tissue accumulation in DIO, and this could mitigate the development of obesity associated lung pathologies.

DIO resulted in a significant reduction in gastrocnemius muscle myocyte size in WTHFD compared to WTCD, and the deletion of *slc7a8* decreased this effect of DIO (KOHFD) on myocytes size (Figure 6G). Additionally, peri-muscular adipose tissue accumulation, which was observed to increase in muscle of WTHFD, decreased in KOHFD (Figure 6E & F). Peri-muscular adipose tissue has previously been shown to promote age and obesity related muscle atrophy by increasing muscle senescence[27]. Hence, a decrease in lipid accumulation due to *slc7a8* deletion in our study suggests an improvement in DIO associated muscular disease. Conversely, there was no significant difference in the gastrocnemius muscle macrophage profile between WTHFD and KOHFD.

The development of cardiovascular diseases is associated with an increase in adiposity[28]. In DIO, the heart of WTHFD showed greater accumulation of epicardial adipose tissue, which was found to decrease in the absence of *slc7a8*, KOHFD (Figure 7). Epicardial adipose tissue is located between the myocardium and epicardium and has properties of brown or beige adipose tissue. It is important for maintaining energy homeostasis and thermoregulation of the heart[29]. However, accumulation of epicardial adipose tissue is associated with increasing BMI and poses a risk for the development of cardiovascular disease[28].

Renal injury and disease have been associated with obesity and studies in mice have documented renal morphological changes due to HFD[30, 31]. An increase in glomerular size and accumulation of lipid droplets in the kidneys was observed in DIO in WTHFD, suggesting an increase in body weight could contribute to renal abnormalities. These changes associated with a greater risk of renal disease was reduced in KOHFD (Figure 9), suggesting that *slc7a8* deletion may improve kidney health in DIO.

This study demonstrates that deletion of *slc7a8* in mice is protective against DIO by significantly reducing adipose tissue mass as well as lipid accumulation in multiple organs and tissues, resulting in improve glucose tolerance in diet induced obesity. Furthermore, our histological findings reveal that the negative effects of DIO on different organs and tissues were improved with *slc7a8* deletion, suggesting a contributing role of this gene to the development of some obesity associated comorbidities. Overall, the results from this study suggest that *slc7a8* could be an important therapeutic target for controlling DIO, as well as for mitigating the development of some of the pathophysiological conditions associated with obesity. Nevertheless, further studies will be required to provide additional knowledge on how *slc7a8* regulates plasma parameters such hormones, lipids and the cytokine inflammatory profile in DIO to reduce lipid accumulation at multiple organs and tissues.

## Materials and methods

### Animals

This study was approved by the Research Ethics Committee, Faculty of Health Sciences and the Animal Ethics Committee, University of Pretoria (Ref. No.: 474/2019). *Slc7a8* (*Slc7a8^tm1Dgen^*) heterozygous and wildtype C57BL/6J mating pairs obtained from Jackson Laboratory (*Bar Harbor*, *Maine*, *United States of America*) were used to generate *Slc7a8* wildtype (WT) and knockout (KO) genotypes. Genotypes were confirmed by PCR (supplementary methods S1). Both WT and KO mice were fed either a high-fat diet (HFD; D12492) or control diet (CD; D12450J) from Research Diets, Inc. (*New Brunswick*, *New Jersey*, *United States of America*) for a period of 14 weeks, with termination time points at weeks 5 and 14. Weekly measurements of weight, food consumption and calorie intake were done. Unless otherwise stated, the nomenclature used for the different genotypes on either a CD or HFD for 14 weeks will be WTCD (wildtype mice on control CD), WTHFD (wildtype mice on HFD), KOCD (*Slc7a8* Knockout mice on control CD) and KOHFD (*Slc7a8* knockout mice on HFD).

### Glucose tolerance and insulin sensitivity tests

Glucose tolerance tests (GTT) and insulin sensitivity tests (IST) were performed in both KO and WT mice prior to introducing them to either CD or HFD. Mice were fasted for 4 hours, and the baseline glucose concentration measured. A 45% D-(+)- glucose solution (*Sigma-Aldrich*, *St. Louis*, *Missouri*, *United States of America*) was then administered interperitoneally at 1.5 mg/g body weight and an insulin solution (*Sigma-Aldrich*, *St. Louis*, *Missouri*, *United States of America*) at 0.8mU/g body weight for GTT and IST, respectively. Blood from the tail vein was used to measure glucose concentration at 15, 30, 60, 90 and 120 minutes using an Accu-Check Instant Blood Glucose Meter (*Roche Diagnostics*, *Basel*, *Switzerland*).

### Histology and immunohistochemistry of mouse tissues and organs

Mice on either CD or HFD were euthanised at week 5 and 14 followed by the collection of white adipose tissue from the inguinal (iWAT), perigonadal (pWAT) and mesenteric (mWAT) depots; interscapular brown adipose tissue (BAT); and liver, kidneys, heart, brain, lungs and gastrocnemius muscle. 10% formalin fixed paraffin embedded (FFPE) tissue sections were processed for histological analysis.

FFPE tissue sections were cut using a microtome and baked at 62°C for 20 minutes followed by haematoxylin and eosin (H&E) staining using a Leica Autostainer XL (*Leica Microsystems*, *Wetzlar*, *Germany*). Slides were mounted with DPX (distyrene, plasticiser, xylene) and imaged using an Axiocam 305 color microscope camera (*ZEISS*, *Oberkochen*, *Germany*) and ZEN 2.6 blue edition software (*ZEISS*).

Immunohistochemical analysis of macrophages was performed as previously described[31]. Briefly, tissue sections were stained with F4/80 rat anti mouse antibody clone A3-1 (*Bio-Rad Laboratories*, *Sandton*, *Johannesburg*, *South Africa*). FFPE sections were baked overnight at 54°C, followed by dewaxing in xylene. The sections were then hydrated through a series of ethanol concentrations, rinsed with distilled water and treated with 3% hydrogen peroxide for 5 minutes at 37°C. Heat-induced epitope retrieval was performed in citrate buffer pH 6,1 (*Dako Target Retrieval Solution S1699*, *Dako*, *Carpinteria*, *California*, *United States of America*) using a 2100 Retriever Unit (*Electron Microscopy Sciences*, *Hatfield*, *Pennsylvania*, *United States of America*). The sections were rinsed in PBS/Tween buffer and treated with 5% Normal Goat Serum (Dako X0907) for 30 minutes after which they were incubated overnight at 4°C with a 1:25 dilution of F4/80 monoclonal rat anti-mouse antibody BM8 (*ThermoFisher Scientific*) or 1:100 F4/80 rat anti mouse antibody clone A3-1 (*Bio-Rad Laboratories*, *Sandton*, *Johannesburg*, *South Africa*). The sections were rinsed in PBS/Tween buffer before incubating for 60 min in 1:200 goat anti-rat IgG (H+L) antibody conjugated to horseradish peroxide (HRP) (Invitrogen, ThermoFisher Scientific). The slides were then developed in 3,3’ diaminobenzidine (DAB) chromogen to visualise F4/80 protein staining. All images were taken and analysed at 20x magnification.

### Statistical and image analyses

Images from H&E and immunohistochemical staining were analysed using ImageJ Fiji (https://imagej.nih.gov/ij/download.html) or Aperio ImageScope version 12.4.3.5008 software (*Leica Biosystems*, *Wetzlar*, *Germany*). Morphometric analysis of the various tissue sections was estimated by measuring the diameter of at least 120 cells distributed across the tissue. Semi-quantitative analysis of F4/80 staining using ImageJ Fiji was done according to the protocol described by Crowe and Jue, 2019[32] to quantify macrophages in the tissues.

Statistical analyses were conducted using GraphPad Prism 5 (*GraphPad Software*, *San Diego*, *California*). Values are expressed as mean ± SEM. One-way ANOVA followed by Bonferroni corrections was used to compare means between three or more categories. When comparing two means, a two-tailed unpaired Student’s t-test was used. Two-way ANOVA with Bonferroni corrections was used where necessary. Statistically significant results are indicated as *P<0.05, **P<0.01, ***P<0.001.

## Acknowledgments

We would like to thank Mr Muchavengwa Chovheya and Mrs Ilse van Rensburg at the Onderstepoort Veterinary Animal Research Unit (OVARU) for assisting with animal care, housing, and experiments; to Mrs Rene Sutherland at the Department of Oral Pathology and Oral Biology for assisting with tissue processing and the Department of Anatomy for allowing us to use their facility including the light microscope.

## Author Contributions

Conceptualization, M.A.A. and M.S.P.; methodology, M.A.A, R.R.P and M.B.vH.; formal analysis, R.R.P and M.A.A.; investigation, R.R.P.; data curation, R.R.P and M.B.vH.; writing—original draft preparation, R.R.P.; writing—review and editing, R.R.P. M.A.A, M.B.vH and M.S.P.; supervision, M.A.A and M.S.P.; project administration, M.A.A.; funding acquisition, M.A.A and M.S.P All authors have read and agreed to the published version of the manuscript.

## Supporting Information

### Supplementary methods

#### S1: Genotyping of mice

Genomic DNA was extracted from tail biopsy of mouse pups using the KAPA Mouse Genotyping Kit (*Wilmington*, *Massachusetts*, *United States of America*) and the KAPA Express Extract Protocol. The extractions were performed in a volume of 100 μl and was set up as follows: 88 μl PCR-grade water, 10 μl of 10X KAPA Extract Express buffer, 2 μl of 1 U/μl KAPA Express Extract enzyme and approximately 2 mm of mouse tail tissue. Enzymatic lysis was performed in the Applied Biosystems 9700 thermal cycler (*Foster City*, *California*, *United States of America*) at 75°C for 10 minutes and enzyme inactivation at 95°C for 5 minutes. The DNA extracts were subsequently diluted 10-fold in 10 mM TRIS-HCL (pH 8.5).

To determine the wildtype, heterozygous and knockout *SLC7A8* genotypes, the following gene-specific primer sequences were used: 5’-CAAATGCCAGCTGTCCTGACCTCAC-3’ forward primer for the wildtype allele, 5’-GGGTGGGATTAGATAAATGCCTGCTCT-3’ forward primer for the knockout allele and 5’-CAGACTTAGGGATGGTGACGCCTAG-3’ for the common reverse primer. All oligonucleotides used in the study were synthesised by Integrated DNA Technologies (*Coralville*, *Iowa*, *United States of America*). The PCR reaction mixture consisted of 6.5 μl of PCR-grade water, 12.5 μl of the KAPA2G Fast Genotyping buffer, 1.25 μl of both the 10 μM wildtype forward primer and 10 μM knockout forward primer, 2.5 μl of 10 μM common reverse primer and 1 μl of the diluted DNA extract. The PCR amplifications were performed in a total volume of 25 μl and cycled in the ABI Applied Biosystems 9700 thermal cycler. The thermal cycling conditions used were as such: 95°C for 3 minutes followed by 95°C for 15 seconds, 60°C for 15 seconds, 72 °C for 15 seconds and a final extension for 2 minutes at 72°C. After amplification, 10 μl of each amplicon was separated on a 2% agarose gel alongside a Thermo Scientific FastRuler Low Range DNA ladder (*Waltham*, *Massachusetts*, *United States of America*). Electrophoresis was performed in 1 x TAE (diluted from UltraPure 10 x TAE buffer (*ThermoFischer Scientific*, *Waltham*, *Massachusetts*, *United States of America*) at 120V for 40 minutes. The gel was stained with Ethidium Bromide Solution, Molecular Grade (*Promega*, *Madison*, *Wisconsin*, *United States of America*) and viewed under UV light using the Molecular Imager Gel Doc XR System (*Bio-Rad*, *Hercules*, *California*, *United States of America*). The expected amplicon sizes were 206bp for the wildtype allele and 390bp for the knockout allele. Only wildtype and knockout mice for the *SLC7A8* gene were used in the study.

#### Supplementary figures

**Figure 1 supplementary:**
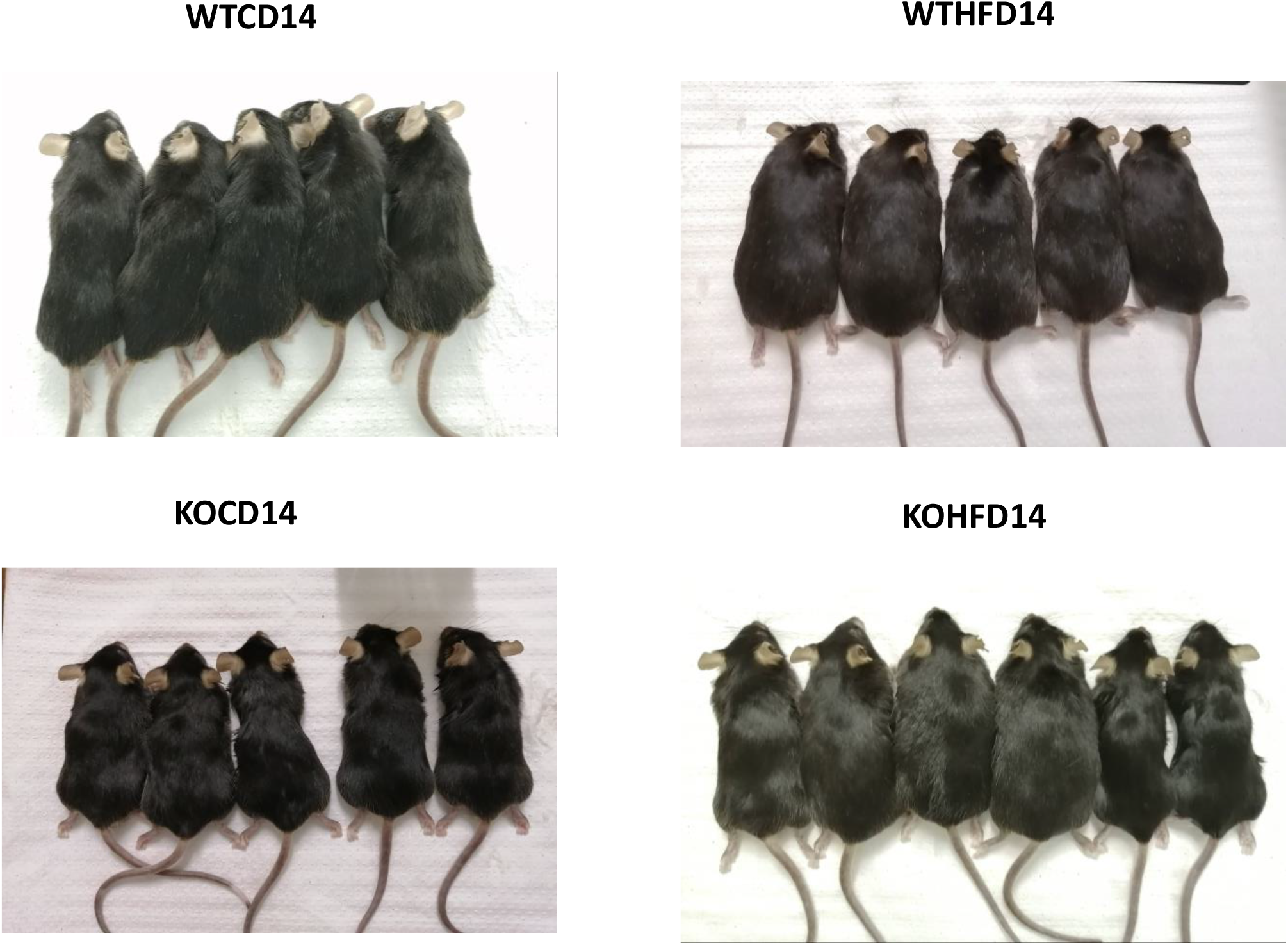
Mice used in the study. The WTHFD14 and KOHFD14 mice look larger in size when compared to WTCD14 and KOCD14, respectively.

**Figure 2 supplementary:**
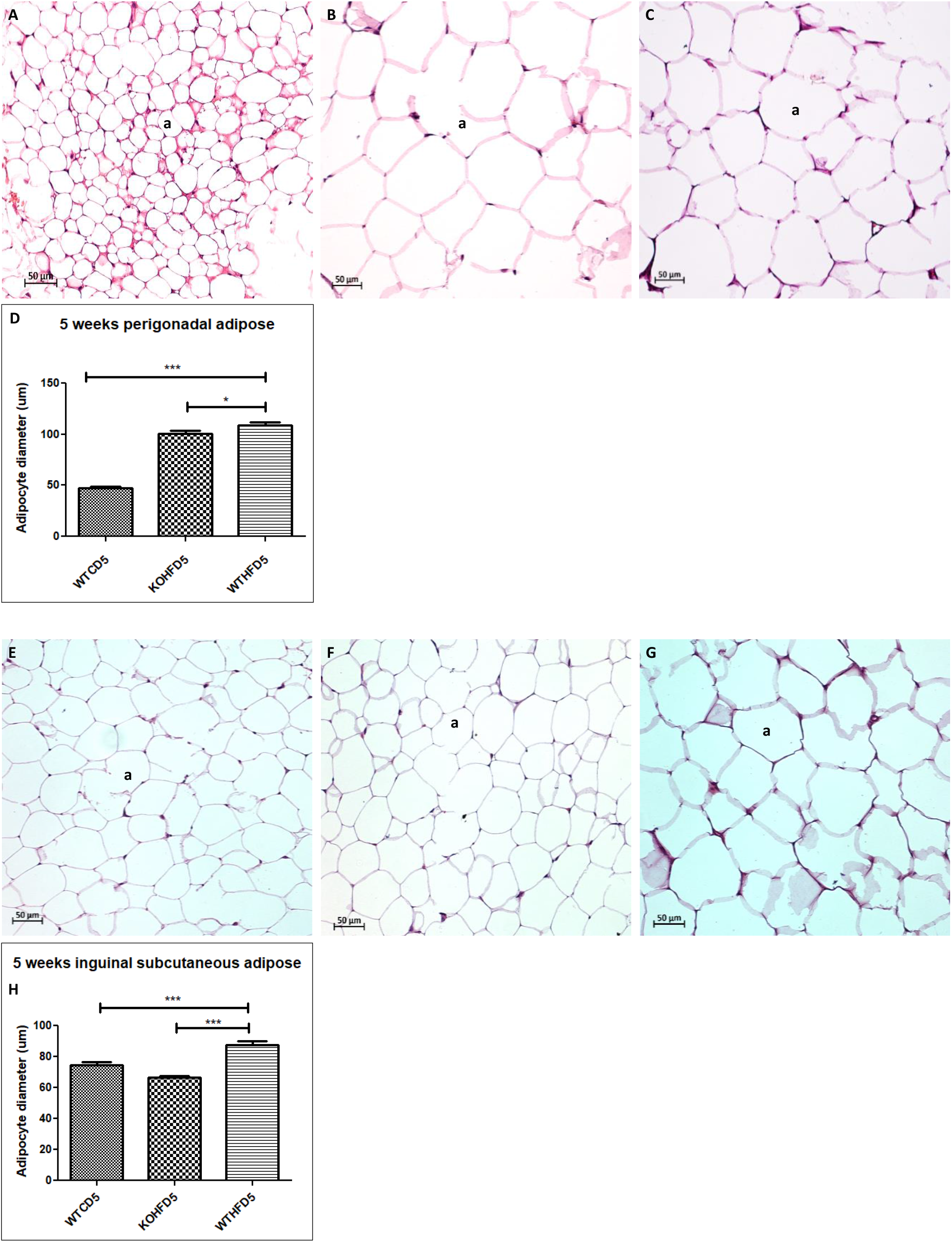
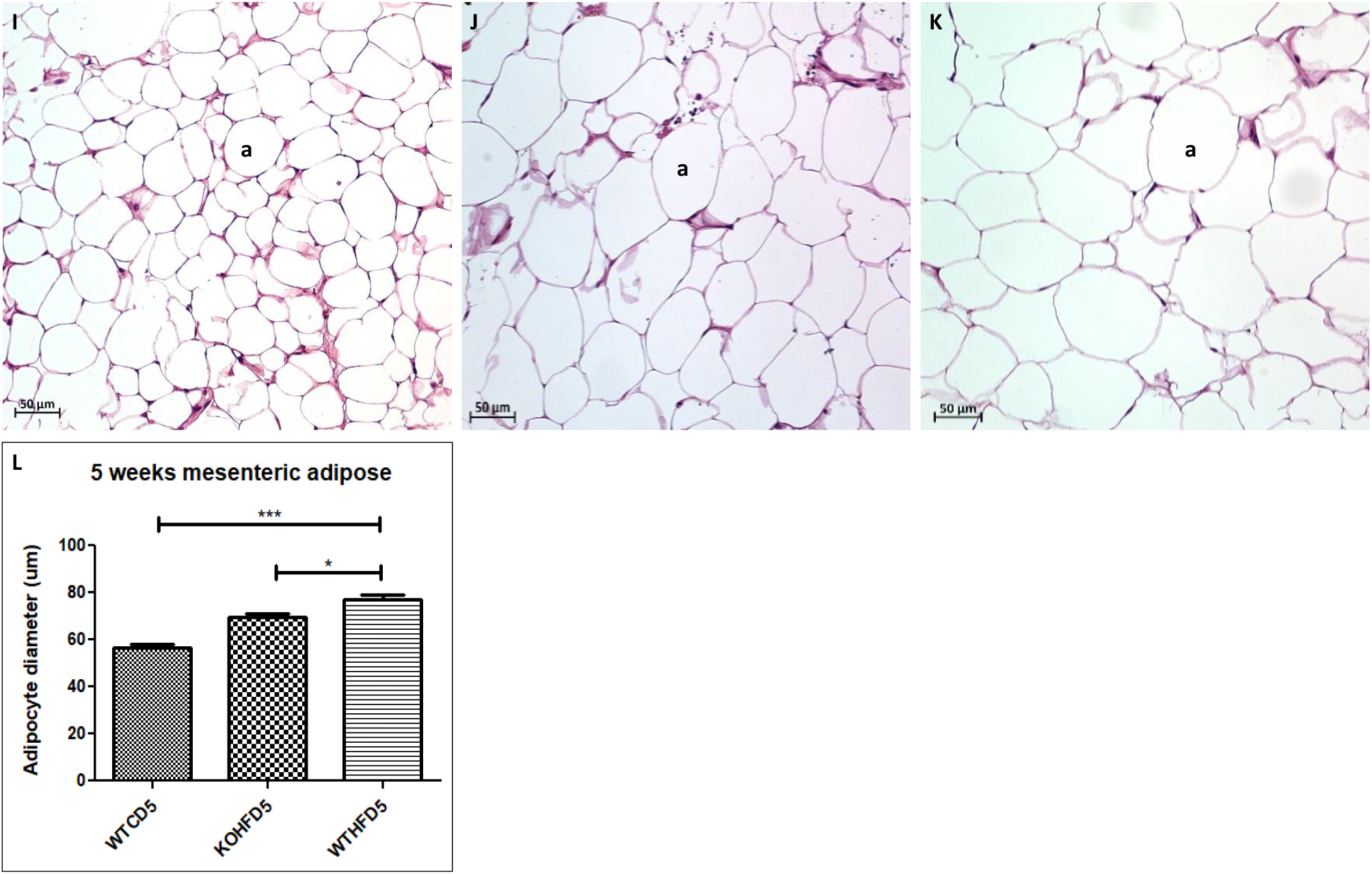
Adipocyte hypertrophy at 5 weeks. Adipocyte diameter of WTHFD, C in pWAT was significantly larger than WTCD, A (p<0.001) and KOHFD, B (p<0.05). In iWAT, WTHFD, G adipocyte hypertrophy was significantly greater (p<0.05) than WTCD, E and KOHFD, F. WTHFD, K in mWAT showed significantly larger adipocytes than WTCD, I (p<0.001) and KOHFD, J (p<0.05). Accumulation of enlarged lipid droplets were observed in WTHFD, O than WTCD, M and KOHFD, N.

**Figure 3 supplementary:**
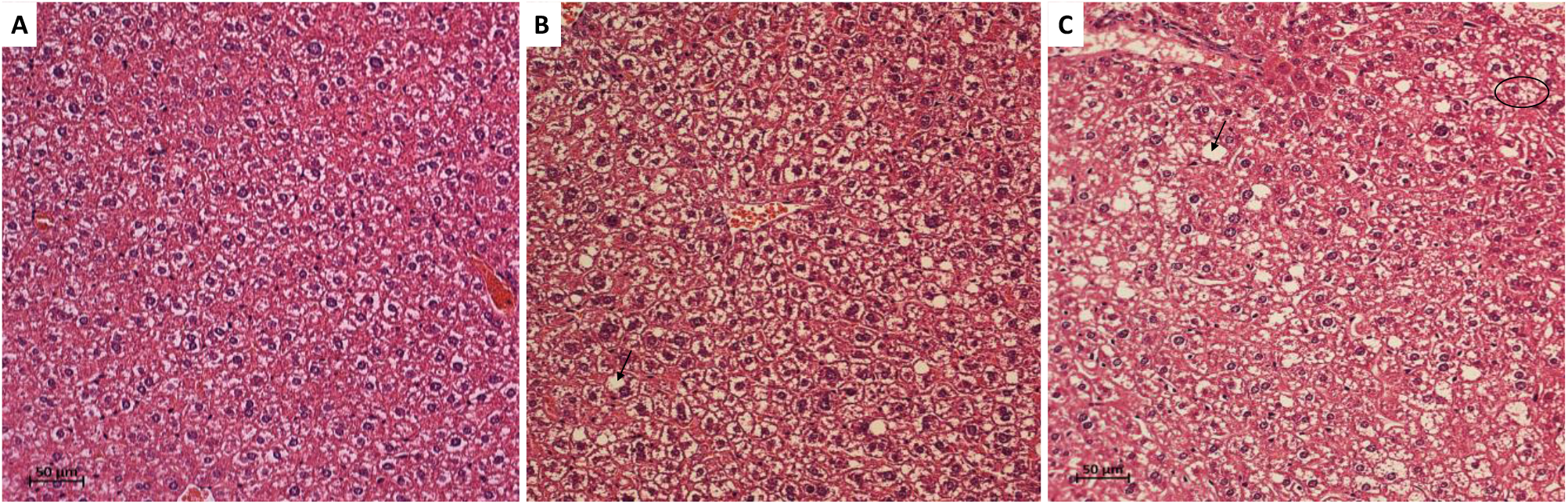
Lipid droplets in the liver at 5 weeks. WTHFD, C and KOHFD, B had lipid droplets in the tissue, while none were observed in WTCD, A.

**Figure 4 supplementary:**
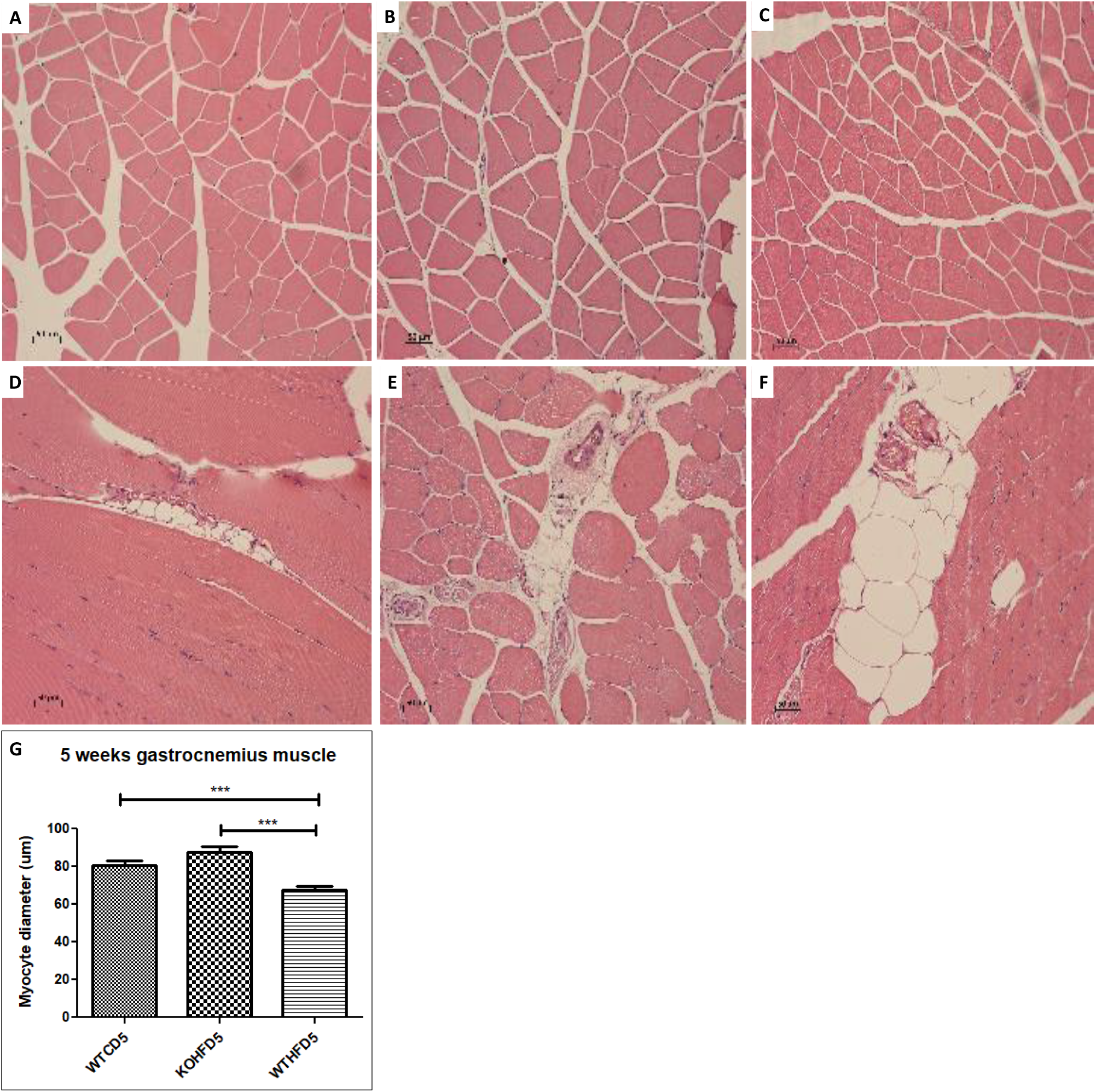
Significantly larger myocytes (p<0.001), G, were observed in the WTCD, A and KOHFD, B in comparison to those in the WTHFD, C group. The distribution of peri-muscular adipose tissue shows that greater accumulation of the adipose was observed in WTHFD, F, compared to WTCD, D and KOHFD, E.

**Figure 5 supplementary:**
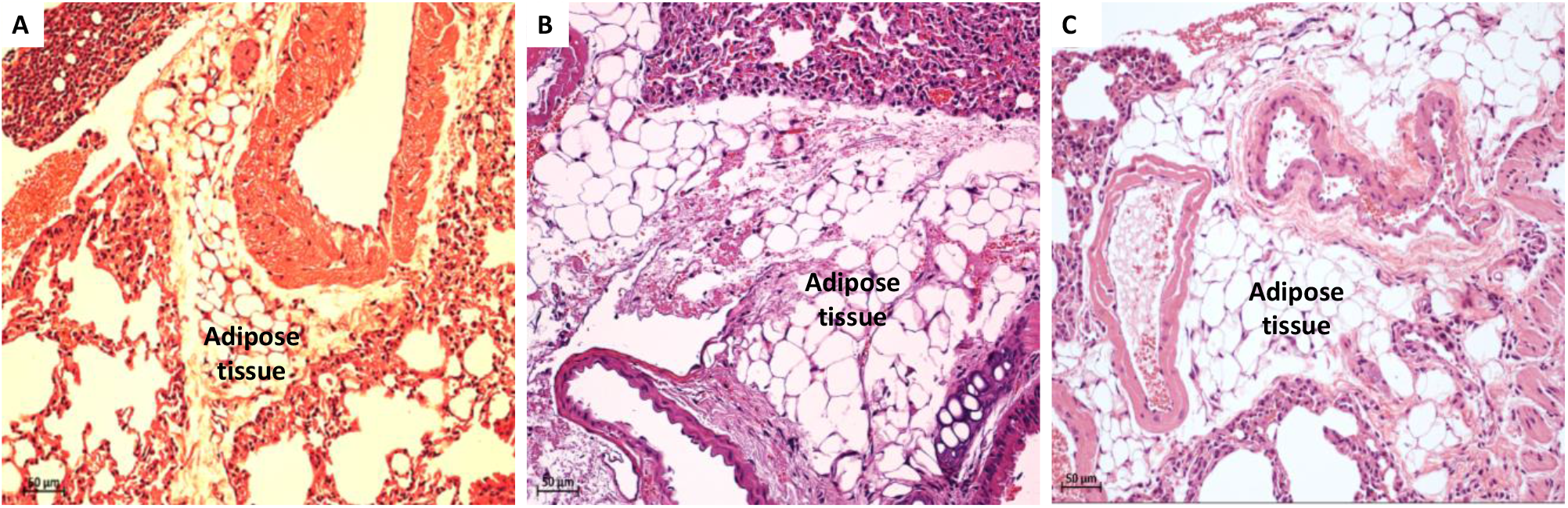
Accumulation of adipose tissue in the lungs. Greater accumulation was observed in WTHFD, C and KOHFD, B in comparison to WTCD, A.

**Figure 6 supplementary:**
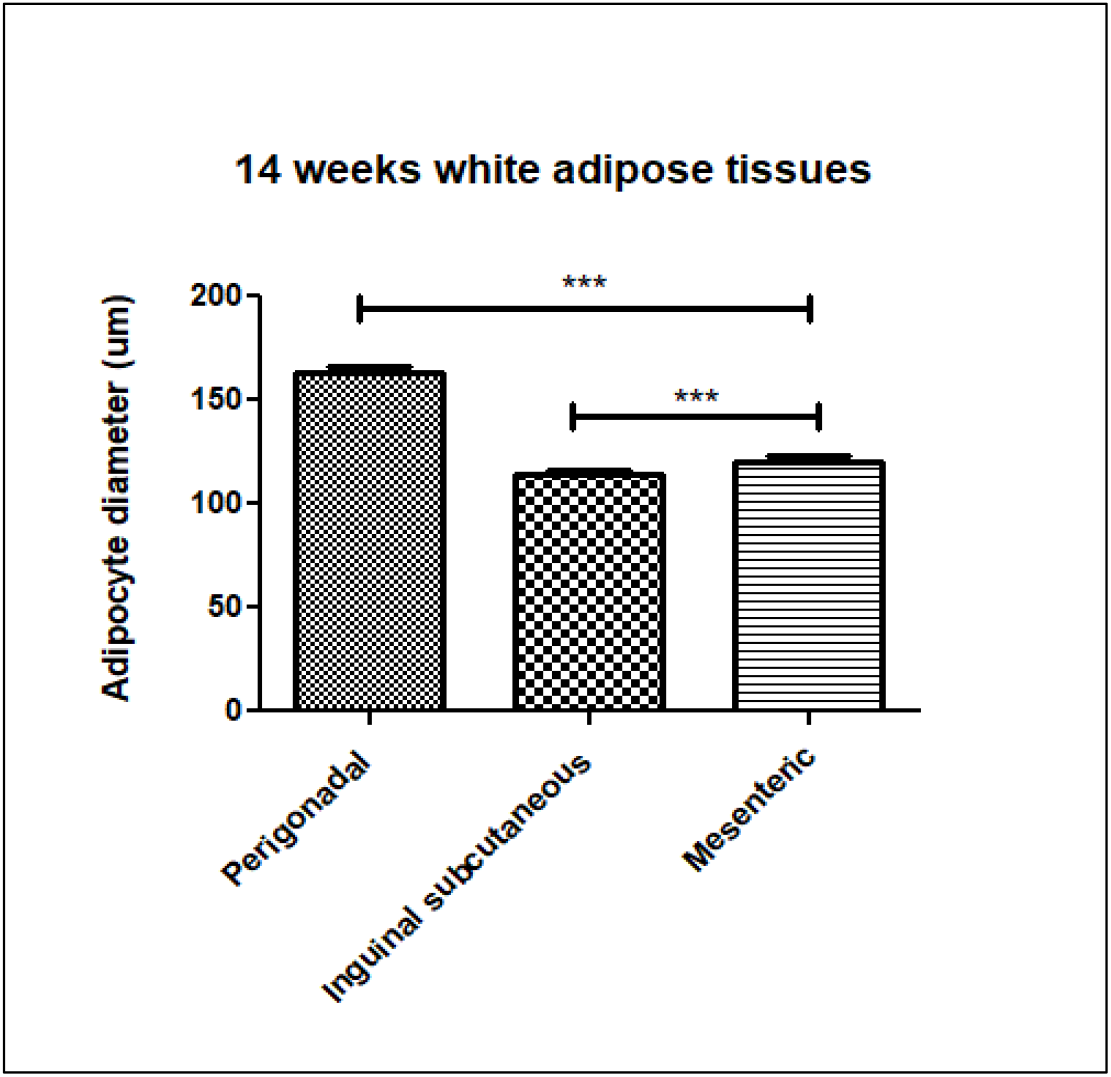
Perigonadal adipose tissue in WTHFD is significantly larger (p<0.001) than inguinal and mesenteric adipose tissues.

